# To be or not to be an anti-CRISPR: AcrIII-1 and the importance of working with native biological systems

**DOI:** 10.1101/2023.01.10.523387

**Authors:** Laura Martínez-Alvarez, Dominic Stickel, Andrea Salegi-Díez, Yuvaraj Bhoobalan-Chitty, Xu Peng

## Abstract

Viral members of the protein family DUF1874 have been reported to act as anti-CRISPR (acr) proteins that degrade cyclic tetra-adenylate (cA_4_), a nucleotide second messenger produced after the activation of several type III CRISPR-Cas systems in bacteria and archaea. Specifically, protein SIRV1 gp29 inhibits type III-A and type III-B CRISPR systems in plasmid-born assays in heterologous systems. In this work, we investigate the function of SIRV1 gp29 and its close homolog SIRV2 gp37 in a native biological context, i.e. in cultures infected by SIRV2. SIRV2 was selected instead of SIRV1 because the latter is not available any more from any laboratories. We demonstrate that gp37 has no anti-CRISPR activity during infection of *Saccharolobus islandicus* LAL14/1 with SIRV2, although it is able to protect SIRV2 from type III targeting when expressed from a plasmid. The inability of gp37 to act as an acr in the native, biological system is due to the protein expression timing: gp37 is a middle/late gene, thus unable to inhibit CRISPR-Cas targeting at the onset of infection. On the other hand, we find that while gp37 is a non-essential gene, it confers a mild replicative advantage to the virus. This advantage is mediated, in hosts with active CRISPR-Cas targeting, by the interaction between gp37 and host protein SiL_1451, which results in the inhibition of the lysine methyltransferase activity of SiL_1451, responsible for extensive methylation of surface lysines of two-thirds of the cellular proteins. Heterologous gene expression from a plasmid-borne non-native promoter has allowed the discovery and characterization of dozens of prokaryotic defense systems in recent years. Although this experimental strategy has several advantages, our study highlights the importance of validating relevant results under native conditions, and the limitations of extrapolating results obtained using heterologous systems.

## INTRODUCTION

Prokaryotes coexist with a range of mobile genetic elements (MGEs, e.g. viruses, conjugative elements) and consequently have developed multiple systems to control or inactivate them^1–4^. Among these, CRISPR-Cas constitutes the only known prokaryotic adaptive immunity. Cas proteins, guided by crRNA that is encoded within CRISPR arrays, mediate the cleavage of complementary foreign nucleic acids^5^. CRISPR-Cas systems are divided in two classes, each containing three types (I, III and IV for class 1 and types II, V, VI for class 2) that are further subdivided into several subtypes^5^. Among the diverse systems, type III are particularly interesting because base-paring of the crRNA with the target RNA elicits specific target degradation by the Cas7-backbone and activates the Cas10 subunit, which in turn degrades DNA non-specifically via its nuclease HD domain^6–8^ and synthesizes cyclic oligoadenylates (cA) via its Palm domain^9–11^. The cyclic oligonucleotide messengers amplify the type III immune response through the activation of CRISPR-Cas associated proteins critical for robust immunity^12–17^, such as Csx1 in *Saccharolobus islandicus* (previously *Sulfolobus islandicus*), a non-specific RNase activated by cyclic tetra-adenylate (cA_4_) produced by its subtype III-B CRISPR-Cas system^15,18^.

Accordingly, viruses possess strategies to counteract cellular defenses and a mechanism to inhibit CRISPR-Cas systems involves anti-CRISPR proteins (Acrs), of which two were reported to work in the evasion of type III CRISPR-Cas targeting. AcrIIIB1 is found in archaeal viruses of the Ligamenvirales order and is required for rudivirus SIRV2 to circumvent *S. islandicus* type III-B immunity through binding to the host type III-B effector complex^19^. The other type III inhibitor, AcrIII-1, is widespread in prokaryotic viruses and its RING nuclease activity rapidly and specifically degrades cA_4,_ thus being able to neutralize type III subtypes that synthesize this nucleotide messenger^20^. Importantly, the activity of the AcrIII-1 family was characterized in heterologous systems using the *acrIII-1* member SIRV1 *gp29* expressed from a plasmid^20^. While rudivirus isolate SIRV1 is no longer available in any laboratories, its closest relative SIRV2 (87.27% average nucleotide identity and thus corresponding to the same viral species) encodes an AcrIII-1 homolog (SIRV2 gp37) with 85% amino acid identity. In this work, we investigate the role of the AcrIII-1 homolog SIRV2 gp37 during SIRV2-infection of its host *S. islandicus* LAL14/1 and demonstrate that gp37 does not exert Acr activity primarily due to having a late expression pattern. However, gp37 is a multi-functional protein capable of interacting with cellular protein lysine methyltransferase and hereby conferring a mild replication advantage unrelated to its RING nuclease activity. Yet, the ability to cleave cA_4_ becomes relevant in the absence of type I-A CRISPR-Cas targeting, implying a physiological role of this second messenger independent to its so far reported function in CRISPR adaptive immunity. These findings can be extrapolated to SIRV1 gp29, as it shares identical genomic neighborhood and regulation as SIRV2 gp37. While plasmid-based heterologous systems have been essential for the discovery and characterization of novel defense mechanisms and their anti-defense counterparts^1–4,21–24^, our results highlight the importance of using native biological systems to confirm the function of newly characterized defense and anti-defense systems.

## MATERIALS AND METHODS

*Strains and growth conditions*. Strains of *S. islandicus* LAL14/1 Δarrays (CRISPR(-) host), Cmr-α (CRISPR III-B(+) host) and Δ*cas6(I-D)* (CRISPR(+) host) (Supplementary Table 1) were grown in basic salts medium^25^ supplemented with 0.2% sucrose, 0.2% casamino acids, vitamin mixture and 0.002% uracil (SCVU medium) at 78°C with agitation. Uracil was removed from the medium composition after plasmid complementation of the Δ*pyrEF* strains. Inducible expression of mini-CRISPR arrays, protein expression or knockdown cassettes was achieved by addition of 0.2% D-arabinose. Strains of *Escherichia coli* NEB Stable (New England Biolabs) or Rossetta (DE3) were used for plasmid cloning and protein expression, respectively and grown in LB medium supplemented with antibiotics. Cellular and viral strains used in this work are listed in Supplementary Table 1.

*Virus propagation and quantification*. Cultures of *S. islandicus* LAL14/1 Δarrays at absorbance at 600 nm (Abs_600_) of 0.2 were infected with the desired virus strain (multiplicity of infection or MOI between 0.01-0.001) and cell turbidity was monitored until a decrease in optical density was observed (around 48 hours post-infection). Cell debris was removed by centrifugation at 10 000 x g, 15 min and virus present in the supernatant was determined by plaque assay as described previously^26^. Two milliliters cells of Abs_600_ 0.2 were mixed with serial dilutions of the supernatant and incubated at 78°C for 30 min, then mixed with 2 mL pre-warmed 0.4% Gelzan (Merck) and immediately plated on pre-warmed SCVU-0.7% gelzan plates. Plates were tightly closed and incubated in a plastic box at 78°C for around two days.

*Average nucleotide identity calculation.* Average nucleotide identity (ANI) between SIRV1 and SIRV2 were calculated using the ANI calculator tool from the enveomics collection (http://enve-omics.ce.gatech.edu/).

*Infection of cell cultures*. Triplicate cultures of the appropriate *S. islandicus* strain were infected at Abs_600_ 0.1 with the corresponding virus at the MOI indicated for each experiment and culture growth curves were determined by measuring the absorbance at 600 nm at regular time-points. One-milliliter samples were taken at the indicated time-points to quantify virus in the supernatant as described above.

*Plasmids, oligonucleotides and DNA sequencing*. Plasmids and oligonucleotides used in this work are listed in Supplementary Tables 2-3. Oligonucleotide synthesis was carried out by Integrated DNA Technologies, Inc. Sanger sequencing used for construct verification was done by Eurofins Genomics.

*Plasmid construction*. Polymerase chain reactions (PCR) for cloning were carried out using Phanta Max Super-Fidelity DNA polymerase (Nanjing Vazyme Biotech Co. Ltd). PCR reactions for colony screening and verification of mutants were done using Taq DNA polymerase (Thermo Scientific). For expression of proteins in *E. coli*, genes were PCR amplified from *S. islandicus* LAL14/1 or SIRV2 genomic DNA with the corresponding primers and inserted in the MCS of vectors p15AIE, pET30 or pETDuet (Supplementary Table 2). For expression of proteins in *S. islandicus*, genes were PCR amplified and cloned into pEXA3^27^. For constructions of virus strains we employed CRISPR-based genetic editing method^26,28^. The corresponding protospacer was cloned into the mini-CRISPR array of the pGE-based vectors^29^ and suitable donor DNA for homologous recombination was inserted at the indicated restriction sites. Knockdown plasmid was constructed by inserting a subtype I-A protospacer targeting SiL_1451 into pGE1. Efficiency of silencing was determined by RT-qPCR and through the determination of protein methylation levels by western blot. Alanine screening mutants of gp37 and SiL_1451 were constructed through site-directed mutagenesis^30^ using the corresponding primers in Supplementary Table 3.

*Cyclic oligoadenylate cleavage assays*. Cyclic oligoadenylate was generated as described previously^9^. Briefly, around 40 nM Cmr-α complex containing only crRNA SS1 targeting the *lacS* gene was mixed with 100 nM target RNA SS1-46, approximately 1 nM α^32^ATP (Perkin Elmer Waltham), 0.1 mM cold ATP, 10 mM Mn^2+^, 5 mM DTT and 20 mM MES pH 6, and incubated at 70°C for 30 min, followed by heat-inactivation at 95°C for 5 min. For single turnover kinetics, 4 μM protein and cOA mix (1:40 ratio) were incubated in 20 mM MES pH 6 and 10 mM Mn^2+^ in 20 uL reactions. Samples were incubated at 70°C for the indicated times and cleavage reaction was stopped by the addition of phenol:chloroform (Thermo Scientific) in a 1:1 ratio. The aqueous phase was recovered and cOA degradation was visualized by electrophoresis in 24% urea-polyacrylamide gels in TBE buffer, followed by phosphor imaging with a Typhoon FLA 700 (GE Healthcare). Kinetic analysis was done as described previously^20^, quantifying cOA_4_ using ImageJ^31^ and fitting the data to a single exponential curve (y=m1 + m2*(1-exp(-m3*x)); m1=0.1, m2=1 and m3=1) using GraphPad Prism version 9.4.1 (681). Raw data for kinetic analysis are available in Supplementary Table 4.

*Construction of SIRV2 strains.* Overnight cultures of *S. islandicus* Δarrays harboring the corresponding genome-editing plasmid were infected with the corresponding parental virus (SIRV2M or SIRV2M Δ*acrIIIB1,* see Supplementary Table 1) at an MOI of 0.1 and incubated overnight. Virus present in the supernatant was collected by centrifugation at 4000 x g for 5 min and 10 μL of the supernatant were used to infect a fresh culture. The process was repeated until the desired virus strain could be detected by PCR (oligonucleotides are listed in Supplementary Table 3). Virus mutants were finally purified through plaque assay.

*Time-course of virus infection*. *S. islandicus* cultures were grown to an optical density of 0.1-0.2 and infected with the corresponding SIRV2 strain at the indicated multiplicity of infection. Growth was monitored to up to 72 h by measuring Abs_600_ and culture samples taken at different time-points to determine virus in the supernatant or for western-blot analysis.

*Protein expression and purification*. Constructs were transformed into *E. coli* Stable cells and plasmid sequence was verified through Sanger sequencing. For protein expression in *E. coli*, plasmids were transformed into Rossetta (DE3) competent cells and 0.5 L of culture were grown in LB medium at 37°C until Abs_600_ reached 0.8, when protein expression was induced by addition of 0.3 mM isopropyl-1-thio-β-D-galactopyranoside (IPTG) and cells were grown at 18°C overnight before harvesting through centrifugation at 10 000 x g for 10 min, resuspended in the appropriate lysis buffer and stored at –80°C until purification. Samples expressing gp37 or gp29 were resuspended in gp37-lysis buffer (50 mM Tris-HCl pH 8, 200 mM NaCl, 5% (v/v) glycerol and 30 mM imidazole), while purification of SiL_1451 and Cren7 was done as described previously^32,33^. For protein expression in *S. islandicus*, plasmids were electroporated into the corresponding strain as described previously^26^. The desired volume of culture (starting Abs_600_ 0.05) was grown at 78°C under agitation and protein expression was induced through the addition of 0.2% D-arabinose to the medium and cells were collected at Abs_600_ around 0.8 by centrifugation, resuspended in the appropriate lysis buffer and stored at –80°C. For lysis of cells, samples were thawed, sonicated (30 cycles of a 3 s pulse plus a 3 s pulse off) and lysed using a French press (FPG 12800, Homogeneising systems LTD). Lysates were clarified of cell debris by two rounds of centrifugation at 12 500 x g for 30 min at 4°C and filtered using a nitrocellulose syringe filter (0.45 μm pore size). His-tagged proteins were purified using HisTrap HP columns (Cytiva). Hemagglutinine-tagged proteins were purified using anti-HA agarose (Thermo Scientific^TM^). Fractions were pooled and concentrated using ultrafiltration units Protein Concentrator PES (3 kDa MWCO) (Pierce^TM^) and loaded onto a Superdex 75 10/300 GL column. Samples of gp37 and gp29 were eluted in gp37-FPLC-buffer (50 mM Tris-HCl pH 8, 200 mM NaCl and 5% (v/v) glycerol). Protein purification was monitored using 12% SDS-PAGE and protein concentration was determined with the Qubit™ protein assay (Thermo Fisher Scientific, catalog no.Q33211).

*Pull-down of N-terminally HA-tagged gp37*. To purify the gene product of *gp37* and any potential interaction partners, 12 L of *S. islandicus* Δarrays were grown to Abs_600_ 0.2 and infected with SIRV2M N-HA-gp37 engineered to express a N-terminal hemagglutinin tagged gp37 (MOI 0.5-1) and incubated for 3 h at 78°C under agitation. Cells were recovered through centrifugation at 10 000 x g for 10 min and the pellet was resuspended in TBS buffer (20 mM Tris HCl pH 7.6 and 150 mM NaCl). Protein was purified using purified anti-HA agarose (Thermo Fisher Scientific) as described above and fractions were analyzed by SDS-PAGE. A fraction containing gp37 and co-purifying protein was analyzed by liquid chromatography-mass spectrometry using the commercial service of Alphalyse, Denmark.

*Western blot and antibodies*. For western blot analysis of uninfected and infected cell extracts, cells from samples of approximately 15 mL were recovered by centrifugation at 10 000 x g, 1 min and resuspended in 2 mL WB lysis buffer (25 mM Tris-HCl pH 7.5, 150 mM NaCl, 1 mM EDTA, 0.1% Triton). Cell lysis was achieved through 15 cycles of sonication (3 s pulse on and 3 s pulse off). Total cell extracts or fractions from protein purification were resolved on 12% SDS-PAGE and transferred to a nitrocellulose membrane using a TransBlot® semi-dry transfer unit (Bio-Rad Laboratories, Inc) and transfer buffer (48 mM Tris base, 39 mM glycine, 10% EtOH, 0.05% Tween 20) at 15 V for 30 min. The membrane was blocked with TBS-0.5% Tween 20 (TBST) buffer with 5 % skimmed milk or 0.5% bovine serum albumin (BSA, for the detection of lysine methylation) for 30 min at room temperature and incubated overnight at 4°C in TBST with 1% skimmed milk or BSA and the corresponding primary antibody. The membrane was washed with TBS, followed by a 1 h incubation at room temperature in TBST-1% skimmed milk/BSA and the corresponding secondary antibody (1:10 000 dilution) and washed with TBS. Finally, the SuperSignal™ West Pico Plus Chemiluminiscent Substrate (Thermo Fisher Scientific) was used for detection via chemiluminiscence. The rabbit anti-gp37 polyclonal antibody was produced by Innovagen AB (Sweden) using recombinant C-terminally His-tagged gp37 expressed in *E. coli* and purified as described above. The rabbit polyclonal antibodies anti-gp17, anti-gp48 and anti-aIF2β were described previously^19,34,35^ and used in a 1:10 000 dilution. The monoclonal HA-epitope tag antibody 2-2.2.14 (Invitrogen, catalog no. 26183) was used for detection of hemagglutinin-tagged proteins (1:5000 dilution). The rabbit polyclonal antibody to methylated lysines (Abcam, catalog no. ab23366) was used for the detection of protein methylation levels (dilution 1:5000). The secondary antibodies goat anti-mouse IgG (H+L) HRP (Thermo Scientific, catalog no. 32430) and goat anti-rabbit IgG-peroxidase (Sigma-Aldrich, catalog no. A0545) were used in a 1:10 000 dilution.

*RNA extraction, cDNA synthesis and real-time quantitative PCR (RT-qPCR)*. Samples of the control strain *S. islandicus* LAL14/1 Δarrays pGE1 and the SiL_1451-knockdown strain *S. islandicus* LAL14/1 Δarrays KD.SiL_1451 were grown in medium supplemented with 0.2% arabinose for approximately 12 h before cell recovery through centrifugation. RNA was extracted using TRI reagent (Sigma-Aldrich, catalog no. 93289) and DNase I-treated (Thermo Fisher Scientific, catalog no. EN0525) according to the instructions of the manufacturers. After verification of RNA integrity through gel electrophoresis, cDNA synthesis was carried out using RevertAid RT reverse transcription kit using random hexamers (Thermo Fisher Scientific, catalog no. K1691), including reactions without reverse transcriptase as controls to discard DNA contamination. RT-qPCR was done using the PowerUP SYBR Green mastermix (Applied Biosystems, catalogue no. A25742) as recommended by the provider for a 10 μL reaction volume and 2 μL diluted cDNA (1:10 dilution). Reactions were amplified on a Quant Studio 3 (Applied Biosystems) with the following program: 50°C for 2 min, 95°C for 2 min and 40 cycles of 95°C for 15 s followed by 58°C for 1 min. Primers used to amplify SiL_1451 and 16S rRNA (used as reference) are described in Supplementary Table 3. The results correspond to the data from three biological samples with three technical replicates each and the SiL_1451 relative level of expression in the knockdown strain KD.SiL_1451 with respect to the control strain was calculated using the threshold cycle (ΔC_T_) method^36^. The C_T_ values of samples and standard curves are reported in Supplementary Table 5.

*Methyltransferase assay*. In vitro methyltransferase activity was measured using the MTase-Glo™ methyltransferase assay (Promega, catalog no. V7601) following the protocol of the manufacturer, with 50 μL reactions carried out in white, opaque flat-bottom OptiPlate-96 microplates (Perkin-Elmer, catalog no. 6005290) at 50°C for 30 min in SiL_1451-MTase buffer (20 mM Tris pH 8, 50 mM NaCl, 1 mM EDTA, 3 mM MgCl_2_ and 0.1 mg/mL BSA) using 10 μM S-adenosyl-L-methionine (SAM), 0.5 μM SiL_1451 purified enzyme and 8 μM recombinant Cren7 as substrate. To test the effect of gp37 over the methyltransferase activity of SiL_1451 the substrate concentration used was 4 μM Cren7 and 4 μM gp37. Incubations with MTase-Glo Reagent and Detection solution were done at room temperature for 30 min, according to the manufacturer instructions. Luminiscence was measured using a FLUOstar® Omega (BMG-Labtech) plate-reading luminometer.

*Structural models*. The structure of the gp37:SiL_1451 complex was predicted using AlphaFold2 ^37^. As gp29 is reported to be a dimer^20^ and SiL_1451 is a monomer^38^, a dimeric conformation for gp37 was used to model the interaction with a monomeric SiL_1451.

*Virus competition assays*. The indicated viruses for each experiment were mixed in a proportion 1:1 and the virus mix was used to infect cultures of the strains of *S. islandicus* Δ*cas6_(I-D)_* (CRISPR(+) host) and *Δcas6_(I-D)_*.KD.SiL_1451 with an MOI of 0.01, or the strains Δarrays (CRISPR(-) host), Δarrays.KD.SiL_1451, Δarrays_OFFspacer (Δarrays pGE1.IASIRV2gp38.sp), Cmr-α targeting (Cmr-α pC*_gp39_*) and I-A targeting (Δarrays pGE1.IASIRV2gp03.noPAM.sp) with an MOI of 0.001, all at an initial Abs*_600_* of 0.1. Cell density was monitored regularly for 2-4 days until cell death was evident by a decrease in absorbance, at which point 1 mL culture samples were taken and centrifuged at 10 000 x g for 1 min to recover the virus present in the supernatant. This corresponds to a single virus transfer step. To start a new transfer, a 10^-4^ (for Δarrays-based strains) or a 10^-3^ (for Δ*cas6(I-D)*-based strains) dilution of the virus supernatant was used to infect a fresh culture of the same strain (initial Abs*_600_* of 0.1) and processed as described before. Between 6-12 transfers were carried out as indicated for each particular experiment. Presence of each virus strain in the supernatant was determined by PCR using Taq polymerase (Thermo Scientific) and 20 μL reactions containing 1 μL of virus supernatant. The oligonucleotides Dgp37.check.F and Dgp37.check.R were used to detect SIRV2M (parental strain) and SIRV2Δgp37 (gp37 knock-out) using the following program: 95° for 1 min, 25 cycles of 95°C for 15 s, 58°C for 15 s and 72°C for 1 min. The oligonucleotides Check.dgp37.F and Check.gp37.R were used for the detection of SIRV2dgp37 (dead gp37), while Check.gp37wt and Check.gp37.R were used for the detection of SIRV2M with the following program: 95°C for 1 min and 25 cycles of 95°C for 15 s, 58°C for 15 s and 72°C for 15 s. The presence/absence of a virus in the supernatant was determined by the presence of a band of the corresponding PCR product visualized with gel electrophoresis.

## RESULTS

### AcrIII-1 homolog gp37 of SIRV2 is unable to inhibit CRISPR-Cas III-B immunity during infection of the host *S. islandicus* LAL14/1

Previous work characterizing members of the AcrIII-1 family showed that the plasmid-borne gp29 encoded by the archaeal rudivirus SIRV1 conferred protection to an *acrIII-1-*null virus SSeV in the presence of CRISPR-Cas III-B immunity^20^. The results by Athukoralage et al. (2020) were of particular interest to us given that a SIRV2 mutant lacking *acrIIIB1* (*gp48*) but carrying *acrIII-1* (*gp37*) was highly sensitive to CRISPR-Cas type III-B interference^19^, suggestive of no AcrIII activity derived from the SIRV2 *acrIII-1*. SIRV2 is closely related to SIRV1 at the core genome region (87.27% average nucleotide identity) and the two viral genomes share high similarity at the amino acid level and genomic organization (Fig. 1A). Moreover, SIRV2 gp37 is 85% identical to SIRV1 gp29 at the amino acid level and the promoter of the operon containing SIRV2 *gp37* is 88.6% identical to the promoter of the corresponding region in SIRV1 (Fig. 1B). To understand the discrepancy of AcrIII-1 activity between the two works, we set on to investigate the role of SIRV2 gp37 during infection. We selected SIRV2 as our model system because the SIRV1 isolate is no longer available in any laboratories.

**Fig. 1.**
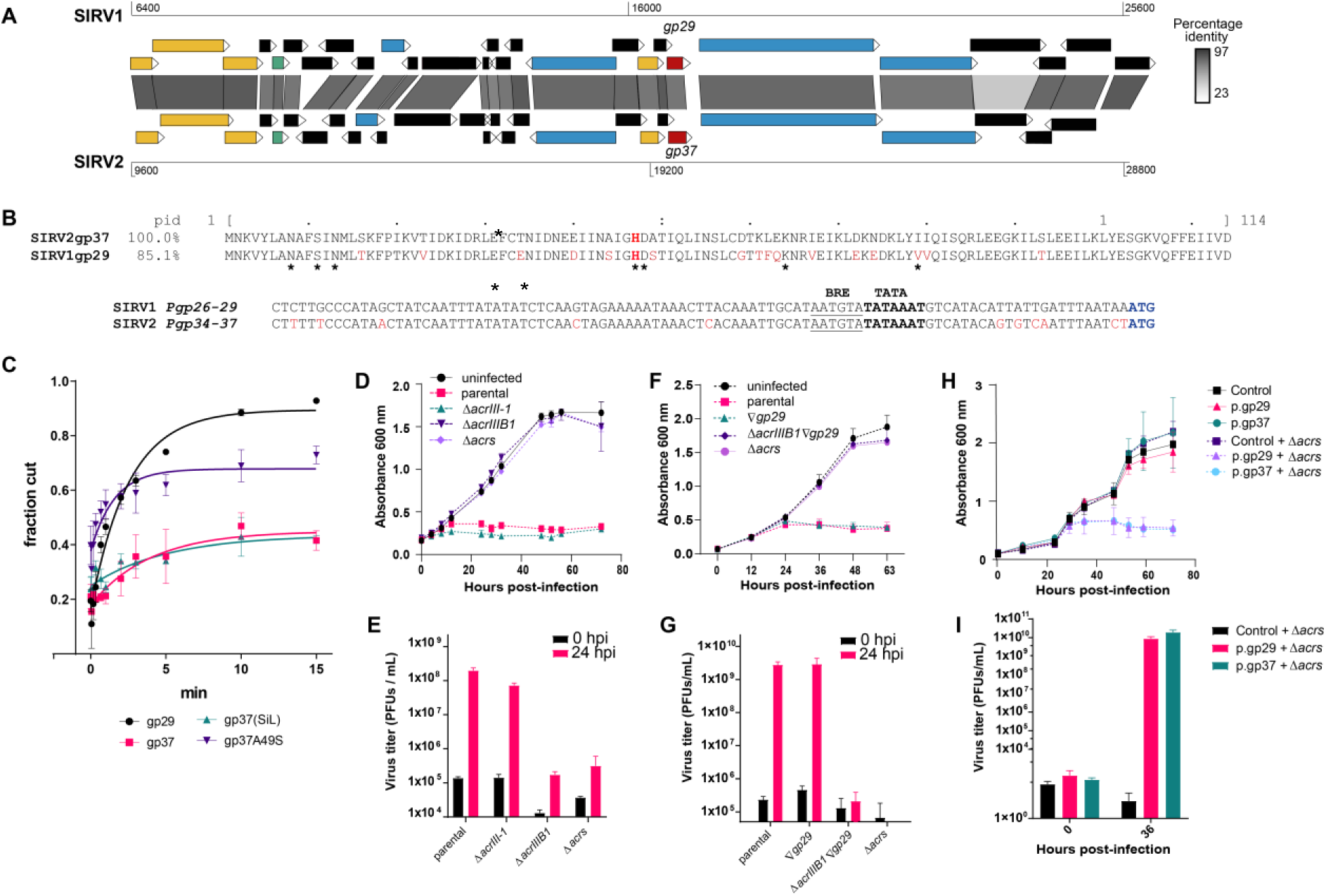
The AcrIII-1 homolog gp37 does not inhibit subtype III-B CRISPR immunity in SIRV2-infected cells. **A**) Comparison of the AcrIII-1 genomic neighborhood in SIRV1 and SIRV2. AcrIII-1 homologs are encoded in a four-gene operon together with highly conserved rudiviral genes. Open-reading frames are colored according to their functional category: yellow – DNA metabolism; green – transcription; blue – virion proteins; red – AcrIII-1 homologs. **B)** Alignment of the promoter region (top) and amino acid sequence (bottom) of AcrIII-1 homologs in SIRV1 and SIRV2. pid – percentage identity. The conserved catalytic histidine is indicated in bold red while residues that differ between the two proteins are shown in red. Nucleotides that differ in *Pgp26-29* with respect to *Pgp34-37* are indicated in red. The start codon is highlighted in blue. The core promoter elements TATA-box and BRE are shown in bold and underlined, respectively. **C)** Kinetic rate of cA4 degradation of the rudiviral AcrIII-1 proteins. Average and s.d. values three biological replicates. Recombinant proteins were expressed and purified from *E. coli*, except gp37 (Sis), expressed and purified from *S. islandicus* LAL14/1. **D-G)** The AcrIII-1 homologs SIRV2 gp37 or SIRV1 gp29 cannot inhibit native targeting by III-B immunity. D,F) CRISPR III-B (+) cultures were infected with the indicated viruses with an MOI of 0.01 and cell growth was monitored by measuring optical density at 600 nm. E,G) Determination of the extracellular virus titer in the cultures from panels D and F. Average and s.d. values from three biological replicates. **H-I)** Plasmid-borne expression of AcrIII-1 homologs exhibits type III-B anti-CRISPR activity. H) Growth curves of CRISPR (+) strains carrying plasmid-borne AcrIII-1 homologs with and without infection by SIRV2 Δacrs. I) Virus titer at 0 and 36 hours post-infection in the supernatant of cultures from panel H. Results from three biological replicates, bars indicate the s.d.

We first compared the ability of the two rudiviral AcrIII-1 homologs to degrade cA_4_ and found that the catalytic rate of SIRV2 gp37 is 50-60% lower than that of SIRV1 gp29 (Fig. 1C). This difference is mainly attributable to a different residue in the catalytic site (a polar residue S49 of SIRV1 gp29 vs. a hydrophobic residue A49 in SIRV2 gp37), as introducing the single change A49S in gp37 increased the catalytic rate of the protein (Fig. 1C).

We then investigated the capacity of AcrIII-1 to protect SIRV2 from type III targeting in its host *Saccharolobus islandicus* LAL14/1. We constructed viral knock-out mutants of *acrIIIB1* (Δ*gp48*) and *acrIII-1* (Δ*gp37*) and assessed their ability to infect *S. islandicus* LAL14/1 Cmr-α pC_gp39_, a strain with active III-B CRISPR targeting. Only viruses encoding *acrIIIB1* were able to lyse the host and replicate to high titer, while the mutant carrying *acrIII-1* alone (no *acrIIIB1*) exhibited little propagation (Fig. 1D-E). To test whether the lack of anti-CRISPR activity of gp37 was due to the lower catalytic activity in comparison to that of SIRV1 *gp29*, we replaced *gp37* with SIRV1 *gp29* (+gp29 strains) in the parental SIRV2 and in the *acrIIIB1* knock-out (Δ*acrIIIB1*∇*gp29*). The presence of *gp29* did not help Δ*acrIIIB1* to overcome type III-B targeting and the replacement of SIRV2 *gp37* by SIRV1 *gp29* did not affect the infectivity of the parental virus (+gp29 vs parental, Fig. 1F-G), demonstrating that the lack of Acr activity in SIRV2-infected cells is not due to differences in the catalytic activity of the homologs.

Athukoralage et al. showed that plasmid-borne SIRV1 gp29 protected virus SSeV from the type III CRISPR defense of *S. islandicus* M.16.4^20^. We reasoned that protein expression from a plasmid might have traits that allow for effective inhibition of type III CRISPR-Cas activity. To test this, we infected cells that express SIRV1 gp29 or SIRV2 gp37 from a plasmid with a virus lacking both *acrIIIB1* and *acrIII-1* (SIRV2 Δacrs). In agreement with the results described by Athukoralage et al., plasmid-borne expression of gp29 homologs, including SIRV2 gp37, is enough to abrogate type III targeting and allows for efficient virus replication (Fig. 1H-I). This result proves that the ring nuclease activity, when expressed from a plasmid, can protect infecting viruses from type III CRISPR-Cas immunity.

### Middle/late expression of the RING nuclease proteins from the viral genome makes them unable to inhibit type III CRISPR-Cas targeting

Rudiviruses exhibit a genomic organization where functionally related genes appear to cluster together, with anti-defense genes located at the termini of the genome and capsid proteins encoded at the middle (Fig. 2A). The RING nuclease gene in SIRV1 and SIRV2 is part of a four-gene operon located in the middle of the genome (Fig. 2A). The other three genes of the operon are part of rudiviral core genome^26^, i.e, genes present in all members, and encode the viral Holliday junction resolvase thought to be involved in genome maturation and packaging^39^. The whole operon exhibits the transcription pattern of middle/late genes^40^. Since the promoter of the operon has a nearly-identical sequence in both viruses (Fig. 1B), we assume that SIRV1 gp29 is also a middle/late gene.

**Figure 2.**
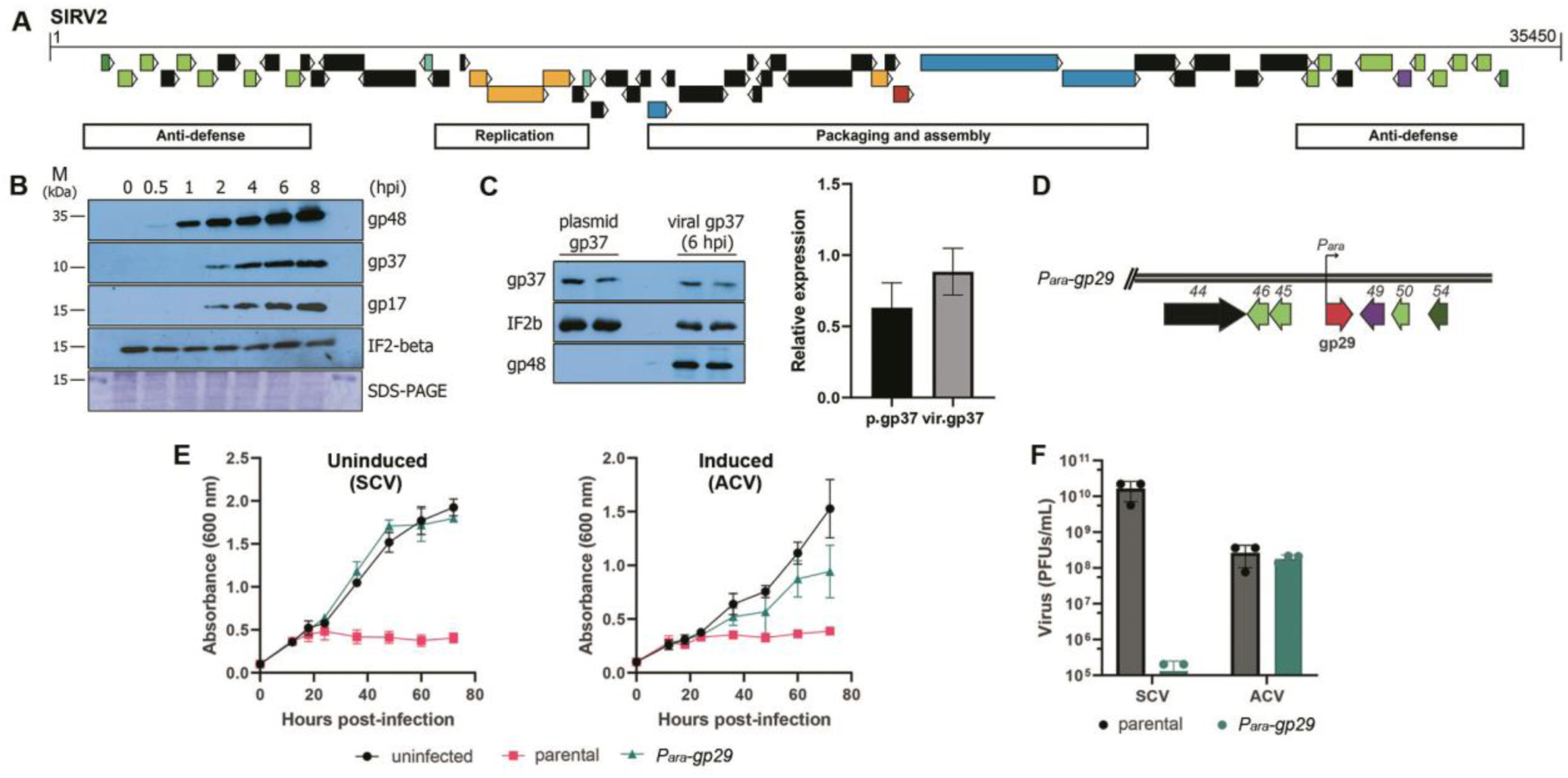
SIRV2 gp37 middle/late expression pattern curbs its ability to exert III-B anti-CRISPR activity. **A**) Functional partitioning of the genome of SIRV2 indicating the regions enriched in genes involved in anti-defense, replication and virion assembly and maturation. The color of the open-reading frames corresponds to: dark green – Aca8 homologs; light green – AcrID1 and AcrIIIB1 homologs; turquoise – transcriptional regulators; yellow – genes involved in DNA replication and metabolism; blue – structural genes; red – AcrIII-1; purple – cell lysis. **B)** Expression kinetics of early (gp48) and middle/late (gp17 and gp37) viral proteins in SIRV2-infected cells determined by western blot. A representative experiment from three biological replicates is shown. **C)** Expression level of plasmid-borne and virally-encoded gp37. Left: representative western blot of plasmidic gp37 and SIRV2-infected lysates at 6 hours post-infection (hpi). Right: average relative levels of gp37 normalized to IF2 expression from three independent biological replicates and bars corresponding to the s.d. **D-F)** Early expression of AcrIII-1 exhibits acr-activity during SIRV2 infection. D) Schematic representation of the genome of the recombinant virus SIRV2 *Para-gp29*, where *acrIIIB1* (*gp48*) is substituted by the *acrIII-1* homolog in SIRV1 under the control of the inducible arabinose promoter (*Para-gp29*). Coloring of open-reading frames is the same as described in panel A. E) Growth curves of CRISPR III-B (+) cultures infected with parental SIRV2 or SIRV2 *Para-gp29* (*P_ara_-gp29*) at an MOI of 0.1 before (left) and after (right) induction of gp29 expression with arabinose (final concentration 0.2%). F) Average and s.d. values of extracellular virus titers from infected cultures at 36 hpi. Data from two biological replicates.

We hypothesized that the absence of type III Acr activity of the native gp37 could be due to its late expression pattern. Western blot of lysates from SIRV2-infected cultures show that, while early gene products such as AcrIIIB1 can be detected as early as 30 minutes post-infection, middle/late gene products like gp37 and gp17 (a ssDNA-binding protein involved in viral genome replication^41^) are detected at 2 hours post-infection (Fig. 2B).

To explore if it is the expression timing or the expression level that impedes gp37 to act as an Acr during SIRV2 infection, we compared the protein level of plasmid-borne gp37 and virus-borne gp37. The expression level of gp37 is, on average, 20% higher in SIRV2-infected cells at 6 hpi than that of plasmid-borne gp37 (Fig. 2C), showing that protein levels during infection would have otherwise been enough to achieve the Acr activity. To further confirm that expression timing is important for the biological function of the viral RING nuclease proteins, we constructed a virus strain lacking *acrIIIB1* and encoding SIRV1 gp29 under the control of an arabinose inducible promoter (SIRV2 *P*_ara_*-gp29*) (Fig. 2D). Without induction, SIRV2 *P*_ara_*-gp29* is unable to replicate (Fig. 2E, left panel; Fig. 2F), but induction of gp29 at the onset of infection recovers the ability of SIRV2 to infect its host (Fig. 2E, right panel; Fig. 2F) and allows SIRV2 *P*_ara_*-gp29* to reach comparable titers to those of the parental virus (Fig. 2F). Taken together, these results demonstrate that early expression of the viral RING nuclease proteins is required for them to exert Acr activity, explaining why SIRV2 gp37, and probably SIRV1 gp29, cannot inhibit type III CRISPR immunity in a natural setting.

### SIRV2 gp37 interacts with and inhibits the host lysine methyltransferase SiL_1451

To investigate the biological function of gp37, which is not a type III CRISPR inhibitor, we looked for interaction partners of the protein during virus infection. We performed a pull-down experiment using an N-terminally HA-tagged gp37 encoded within the viral genome, thus the assay was performed in SIRV2-infected cultures where expression regulation of gp37 follows the native biological conditions. The recombinant SIRV2 HA-gp37 had similar infectivity to the parental strain (Fig. S1), demonstrating that the tag does not affect the infectivity of the virus. Following pull-down a single additional band over the 15 kDa marker position was detected by SDS-PAGE and mass spectrometry showed it to match the 18 kDa host protein lysine-methyltransferase SiL_1451 (Fig. 3A and Fig. S2). The interaction between the two proteins was confirmed with a complementary pull-down assay performed using a C-terminally tagged SiL_1451 expressed from a plasmid, which recovered gp37 from SIRV2-infected cells (Fig. 3B). The yield and the purity of SiL_1451-HA are very low despite arabinose induction of gene transcription, and protein identity was thus confirmed using western blot (Fig. 3B, bottom).

**Figure 3.**
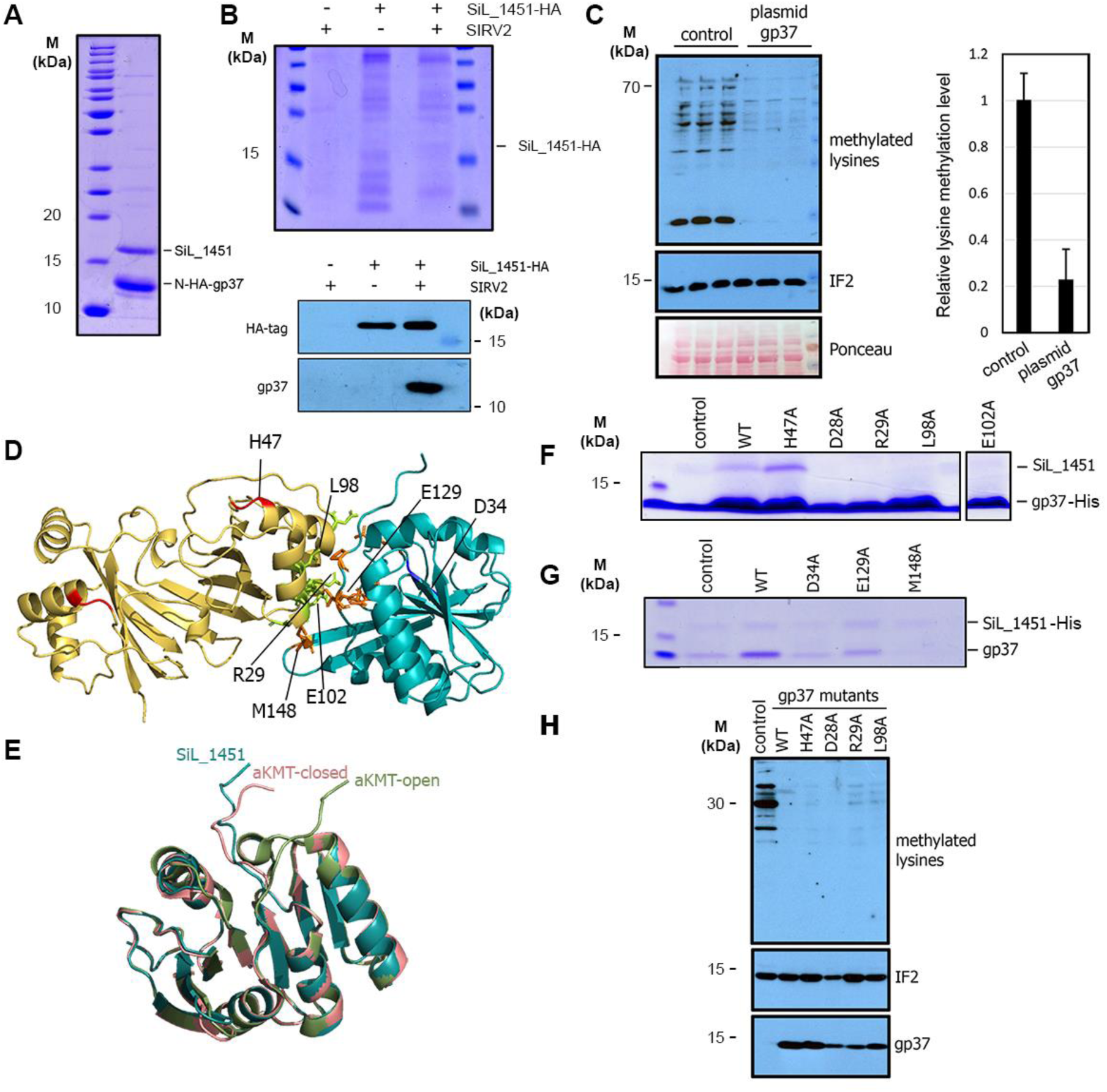
SIRV2 gp37 binds and inhibits the cellular protein lysine methyltransferase SiL_1451. **A-B**) Co-purification of HA-gp37 and SiL_1451. Pull-down assay using a virally encoded gp37 bearing an N-terminal HA-tag **(A)** Pull-down assay with plasmid-borne C-terminally HA-tagged SiL_1451 with or without infection with SIRV2 **(B, top)** and western blot using anti-HA and anti-gp37 antibodies **(B, bottom)**. **C)** Plasmid-expressed gp37 inhibits protein lysine-methylation levels. Left: representative western blot of cellular lysates (*n*=3) using polyclonal antibodies to mono– and di-methyl lysines or translation initiation factor 2 beta (IF2) and total protein stained with Ponceau as a loading controls. Right: average relative lysine methylation levels with s.d. from three biological replicates. **D)** Model of the complex between a gp37 dimer and SiL_1451. Red: residues important for ring nuclease activity. Blue: SiL_1451 catalytic residues. Green and orange: residues involved in the interaction between gp37 and SiL_1451, respectively. **E)** Comparison of the structures of SiL_1451-SAH (aKMT-open,green), SiL_1451-SAM (aKMT-closed, pink) and gp37-bound SiL_1451 (blue). **F)** Pull-down assay of plasmid-expressed C-terminally His-tagged gp37 mutants. **G)** Pull-down of plasmid-expressed untagged gp37 and N-terminally His-tagged SiL_1451 variants. **H)** Representative western blot (*n* = 3) of the lysine methylation levels in cellular lysates from panel G. Translation initiation factor 2 beta (IF2) is used as loading control and SIRV2 gp37 as control of viral infection.

SiL_1451 is known to mediate extensive protein lysine methylation with broad substrate specificity, being responsible for the mono-methylation of one-third of all proteins encoded by *S. islandicus,* including several of the most abundantly expressed proteins^32,42^. Post-translational lysine methylation is widespread in crenarchaeota, albeit the functional significance of this modification is yet to be elucidated^38^. To investigate if the interaction between gp37 and SiL_1451 affects the enzymatic activity of the latter, we determined the levels of intracellular protein methylation in cells carrying a plasmid-borne gp37 by western blot using a polyclonal antibody recognizing mono– and di-methylated lysines. We observed a drastic reduction of 80% in protein methylation levels after induction of gp37, indicating that gp37 inhibits SiL_1451 (Fig. 3C).

To get further insight into the mechanism by which gp37 inhibits the methyltransferase activity of SiL_1451, a model of the interacting proteins was first predicted by Alphafold2 (Fig. 3D). Several contacts are predicted between one chain of the gp37 dimer and the N-terminal and C-terminal ends of SiL_1451, including ionic interactions between D28, R29 and D59 in gp37, and L142, E129 and R133 in SiL_1451, as well as hydrophobic and polar interactions mediated by residues L30, L98, Y101 and E102 in gp37 and P8, V10, P102, L104 and M148 of SiL_1451 (Fig. 3D). A comparison of the conformation of SiL_1451 in our model to the crystal structures of the methyltransferase in the open (cofactor-binding SiL_1451-SAM) and closed (reaction product-binding SiL_1451-SAH) conformations suggests that the interaction with gp37 positions SiL_1451 in a closed conformation where access to the methyltransferase active site is blocked by the placement of the N-terminal portion of the protein (Fig. 3E). If gp37 locks SiL_1451 into an inactive conformation, this could explain the lack of methyltransferase activity observed in the presence of gp37 (Fig. 3C).

The gp37-SiL_1451 model was tested through an alanine-scanning of the presumed interacting residues. We focused on the residues putatively mediating the contact between the two proteins while maintaining their respective catalytic activities. For this, we determined the ring nuclease activity and methyltransferase activity of gp37 and SiL_1451, respectively, and selected the mutants exhibiting activity comparable to the wild-type proteins (Fig. S3A-C). We included as controls the catalytically inactive gp37 H47A and SiL_1451 D34A, previously shown to abrogate cA4 cleavage and lysine methyltransferase activity, respectively^20,38^. Mutation of the gp37 residues D28, R29, L98 and E102 did not impair cA4 degradation, and only M148 mutation in SiL_1451 maintained wild-type-like lysine methylation capacity, while K104 and E129 show halved enzymatic activity (Fig. S3A-C).

Next, we evaluated the interaction capacity of the gp37 mutants through pull-down assays in *S. islandicus* cells expressing plasmid-borne gp37 with a histidine tag attached to the C-terminus. The wild-type and catalytically dead gp37 positive controls co-purify together with SiL_1451, as indicated by the presence of a 18 kDa band corresponding to the lysine methyltransferase (Fig. 3F). This band is missing in all the alanine-mutants tested, indicating an impaired interaction with SiL_1451 (Fig. 3F and Fig. S3D). To test the gp37-interaction capability of the SiL_1451 mutants, pull-down assays were carried out with cells expressing an N-terminally His-tagged SiL_1451 and an untagged gp37 from a single plasmid. Although SiL_1451 has similar low yields as previously seen (Fig. 3B), the wild-type protein co-purifies with gp37, while the catalytically dead mutant D34 and mutant E129 appear both to have a decreased interaction with gp37, and mutant M148 shows the greatest reduction in gp37 co-purification, as the 14 kDa band corresponding to gp37 is barely visible (Fig. 3G). Overall, the results demonstrate that all the tested residues are involved in the physical interaction between the two proteins. Interestingly, a decrease in lysine methylation levels was detected in the lysates of all the gp37 mutants tested in these assays, comparable to the level of methylation in cells expressing the wild-type proteins (Fig. 3H). These results suggest that a single mutation of the residues mediating the gp37-SiL_1451 binding may not be enough to completely abrogate the interaction between gp37 and SiL_1451. An alternative explanation is that the low protein level of SiL_1451 is readily inhibited by the high intracellular level of the plasmid-borne gp37, even if the interaction between the two is impaired.

### SIRV2 gp37 is a multi-functional protein that confers a mild replicative advantage through different mechanisms

In *Sulfolobus* carrying CRISPR type III-B immunity the viral mutant Δ*gp37* exhibited a reduction of around 3-fold in virus yield when compared to the parental virus (Fig. 1D). Moreover, this difference was maintained during infection of a strain without CRISPR-Cas defense (Fig. 4A), suggesting that gp37 confers a mild replicative advantage which is independent of the CRISPR immunity.

**Figure 4.**
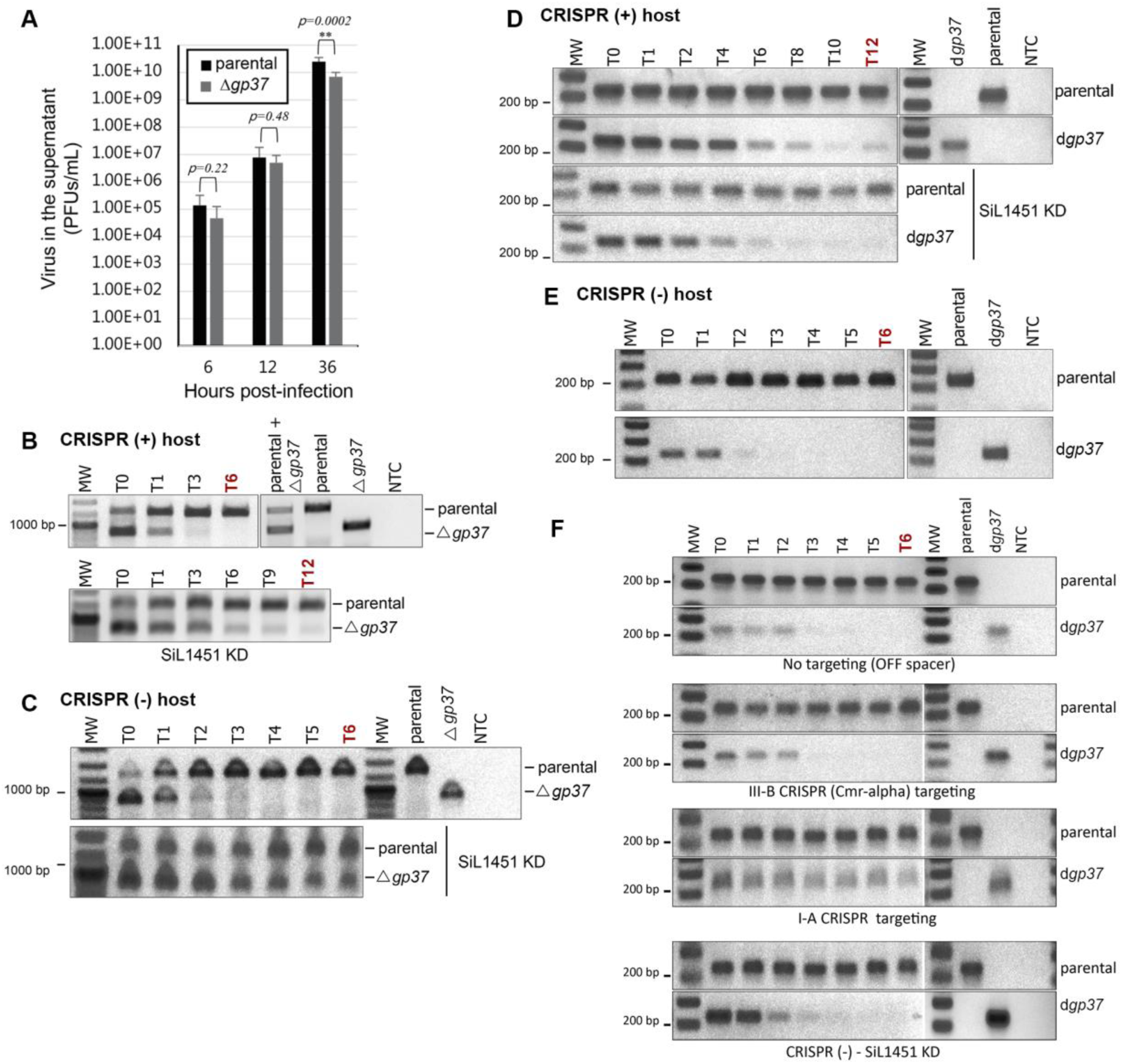
The two activities of SIRV2 gp37 confer viral replicative advantage under different scenarios. **A**) Production of virus progeny (PFUs/mL) in a CRISPR (-) host infected with the parental SIRV2 or a Δgp37 deletion mutant. Data corresponds to the average and s.d. of eight biological replicates. **B-E)** Competition assay between parental SIRV2 (encoding gp37 wt) and a virus lacking gp37 (Δgp37) or encoding a variant of gp37 without ring nuclease activity (dgp37). Presence of viruses in the supernatant was monitored by qualitative PCR using primers specific for each viral strain. Cultures of CRISPR (+) (B-C), CRISPR (+) – *SiL_1451* knockdown (B-C) or CRISPR (-) (D-E) hosts were infected with an equimolar mixture of the indicated viruses at an MOI of approximately 0.0001. **F)** Same as B-E but host cultures correspond to a strain carrying a spacer with no match on SIRV2 (first panel, off spacer), CRISPR (+) III-B cells (second panel), a CRISPR (+) I-A carrying a spacer able to elicit I-A immunity but unable to activate III-B immunity as the crRNA is not complementary to the target gene transcript (third panel), or a CRISPR (-) – *SiL_1451* knockdown host (bottom panel).

To explore the roles of the ring nuclease activity of gp37 and its interaction with SiL_1451 lysine methyltransferase in the gp37-mediated replicative advantage, we constructed the virus strain SIRV2 d*gp37* that encodes the inactive ring nuclease variant gp37 H47A, and confirmed that the expression level of dead gp37 is comparable to that of the wild-type protein (Fig. S4). Additionally, we constructed *SiL_1451* knockdown strains of *S. islandicus* achieving a 60% reduction of the *SiL_1451* mRNA and a decrease of ∼ 50% in the intracellular lysine methylation level (Fig. S5).

Next, we performed competition assays between the parental and the recombinant SIRV2 strains Δ*gp37* (without gp37) or d*gp37* (with abrogated cA_4_ cleavage activity). Viruses were mixed in a 1:1 ratio and used to infect different *S. islandicus* strains at low multiplicity of infection (0.001-0.0001). When complete lysis of the culture was achieved, culture supernatants containing extracellular virus were used to infect fresh cultures for a “transfer” step. This process was repeated for 6-12 transfers and the presence of each virus strain was assessed using qualitative PCR.

In both cells with active CRISPR-Cas targeting or no targeting, SIRV2 Δ*gp37* is quickly out-competed by the parental strain after 2-3 transfers (Fig. 4B and 4C, top panels). However, the disadvantage of lacking gp37 is sensibly neutralized after silencing of SiL_1451, where SIRV2 Δ*gp37* is still detectable in the supernatant after twelve transfers (Fig. 4B and 4C, bottom panels). This suggests that gp37 confers a replicative advantage primarily through its ability to inhibit SiL_1451 and this advantage is independent of CRISPR immunity.

We then explored the role of ring nuclease activity during infection by the competing parental and d*gp37* viruses. The advantage of the parental virus was minimal in cells carrying CRISPR I-A and III-B immunities, with d*gp37* persisting after twelve transfers (Fig. 4D).

However, the ring nuclease activity is critical to the gp37-mediated replicative advantage in cells carrying no CRISPR immunity, as SIRV2 d*gp37* rapidly disappeared from the population after two transfers (Fig. 4E), contrary to what occurred in cells with CRISPR immunity (Fig. 4D). The CRISPR-null strain lacks any CRISPR arrays, but contains fully functional subtype I-A and III-B *cas* gene cassettes^43^. We then selectively activated each of the CRISPR-Cas subtypes by providing a spacer for their specific activation. Virus d*gp37* was out-competed quickly by the parental virus in the control strain carrying a spacer not matching the viral genome, as well as in the strain with subtype III-B targeting (Fig. 4F, first and second panels). However, dgp37 disadvantage was abated in the strain with active I-A targeting (Fig. 4F, third panel), signifying that I-A CRISPR-Cas targeting reduces the need for cA_4_ cleavage or somehow inhibits the ring nuclease activity.

In both host strains with and without CRISPR immunity, *SiL_1451* knockdown does not alter the phenotype of the competition between the parental and the d*gp37* viruses (Fig. 4D and 4F, bottom panels), in contrast to what was observed in the competition assays between the parental and Δ*gp37* viruses. The results imply that gp37 confers mild replicative advantage through at least two pathways. The first inhibits the lysine methyltransferase activity and does not involve the ring nuclease activity nor the host CRISPR immunity. The second involves the ring nuclease activity whose effect is minimized in the presence of type I-A CRISPR targeting.

### Other RING nuclease proteins unlikely act as type III CRISPR inhibitors

Although our results clearly demonstrate that SIRV2 gp37 does not have Acr function in its biological context, they show it can potentially act as such under artificial expression conditions. Thus, we set up to explore if other viral RING nucleases in archaea may exhibit Acr activity by analyzing their genomic context and regulatory sequence. We have recently shown that anti-defense genes in crenarchaeal viruses, including anti-CRISPR proteins, are transcribed from the same promoter driving strong and early expression of the genes, named anti-defense regulatory region (*P_ADE_*) (Bhoobalan-Chitty et al., unpublished). Out of 24 viral RING nuclease genes in crenarchaeal viruses, only two are under the control of *P_ADE_* and located in an anti-defense island a 0.13 ratio, in contrast to AcrID and AcrIIIB1 homologs that have a ratio of 0.54 and 0.75, respectively. An analysis of the genomic neighborhood reveals that seven of the genes are associated to regions enriched in anti-defense genes (AFV3-8, SSV13, SSV.M14.15 and SRV), seven are not associated to anti-defense islands (SIRV1-3, ARV1, AFV9, SIFV1-2) and for the remaining 10 it is not possible to determine the functional association of the neighboring genes (turriviruses, bicaudaviruses) due to lack of information about the protein function and genome organization. Importantly, four of the seven anti-defense loci encoding RING nucleases also encode a recognizable homolog of AcrIIIB1 (Fig. S6), reducing their likelihood of being an AcrIII. While this analysis do not strongly support an Acr role for viral RING nucleases, it suggests that some of these could be involved in anti-defense roles. However, the frequent presence of viral RING nuclease homologs in non-anti-defense genomic contexts together with our results support a role of these proteins beyond inhibition of type III immunity.

As SiL_1451 homologs are restricted to crenarchaea, the interaction between the gp37 and lysine methyltransferase homologs is only relevant for members of this phylum. Importantly, while residues involved in cA_4_ cleavage are highly conserved in vRING nucleases, residues involved in binding of SiL_1451 are more variable and conserved only in a few related rudiviruses (Fig. S7), hence, protein sequence analysis do not support a widespread conservation of the interaction. On the other hand, the lysine methyltransferase homologs are conserved in the crenarchaeota (67-100% amino acid identity), including near-absolute conservation of the identified gp37-interacting residues (Fig. S7). Modelling of complex formation between RING nuclease proteins from crenarchaeal viruses and their cognate host lysine methyltransferase suggests all of the tested protein pairs interact using equivalent residues for binding (Fig. S8). Moreover, complex formation was predicted between SiL_1451 and bacterial RING nucleases YddF from *B. subtilis* and *Listeria boonae*, or the DNA-binding protein Sso7d and the clamp protein PCNA, all known substrates of SiL_1451^32^ (Fig. S8). However, no complex formation was predicted between SiL_1451 and Cren7, another substrate of the crenarchaeal protein methyltransferase. Thus, it is possible the interaction between SiL_1451 and viral RING nucleases is a by-product of the broad substrate recognition of the crenarchaeal lysine methyltransferase that happened to be beneficial for viral replication.

## DISCUSSION

In this study, we demonstrate RING nuclease gp29 from SIRV1 and its homolog gp37 from SIRV2, previously reported to act as type III anti-CRISPR proteins^20^, are unable to inhibit type III targeting in their native biological conditions. We show that lack of Acr activity is due to the late expression timing of the protein during infection and additionally uncover the interaction between gp37 and SiL_1451, a cellular protein lysine methyltransferase, which results in a mild viral replicative advantage that is independent of the RING nuclease activity of gp37.

While gp29 and gp37 are unable to inhibit type III-B immunity when expressed from their native promoter in the viral genome, plasmid-borne proteins protect an *acrIII*-lacking SIRV2 mutant from type III targeting (Fig. 1D-G). This highlights the importance of immediate inhibition of CRISPR-Cas targeting for Acr function and for the establishment of viral infection. The absence of type III Acr activity of gp37 is further supported by the virus competition assays that show cA_4_ cleavage is not required for the gp37-conferred replicative advantage and that in wild-type cells gp37 assists viral infection through the inhibition of SiL_1451 (Fig. 4B-C). Furthermore, I-A CRISPR-Cas targeting is required for the gp37-phenotype, as viral competition in a mutant strain lacking I-A targeting uncovers the benefit of gp37 RING nuclease activity (Fig. 4F). Two findings are inferred from this result: i) first, in the native host the presence of CRISPR-Cas immunity determines the mechanism of action of gp37 through SiL_1451 inhibition; ii) second, cA_4_ degradation is important in the absence of CRISPR-Cas systems, demonstrating the existence of other pathways of cA_4_ generation besides type III systems and a role for this secondary messenger independent of CRISPR immunity. Figure 5 summarizes the above findings and provides a model to explain the role of gp37 alternative activities under different scenarios. The presence of alternative cA_4_ synthesis pathways has especial significance given that nucleotide second messengers are implicated in signal transduction of developmental, physiological and defense processes in all organisms. In particular, our knowledge about the diversity of nucleotide second messengers involved in defense mechanisms in bacteria (e.g. CBASS, cGAS-STING, Pycsar, type III CRISPR-Cas) has greatly expanded in the last years^44–49^. Contrastingly, little is known about nucleotide second messengers in archaea. A recent study identified the presence of several cyclic and linear nucleotide molecules in *S. acidocaldarius* and *H. volcanii,* although their biological role remains to be determined^50^. Up to date, the only described role for cA_4_ is related to type III CRISPR-Cas immunity and future work shall elucidate alternative functions of cA_4_ and other second messengers in archaea, a very likely scenario given that the role of nucleotide messengers in defense is a conserved feature in the other domains of life. We acknowledge our results do not absolutely discard a role as Acr for viral RING nucleases, albeit they introduce the reasonable presumption that these proteins frequently have alternative roles during viral infection. This inference is supported by the analysis of the genomic neighborhood of the RING nuclease genes in archaeal viruses, where around a two thirds of the gp29 homologs are not encoded in recognizable anti-defense islands, as would be expected for *acr*s (Fig. S6). Moreover, we speculate that the interaction between viral RING nucleases and crenarchaeal lysine methyltransferases is the result of the lack of substrate specificity of the latter whose inhibition is beneficial for viral infection of crenarchaeal hosts. This does not rule out a role for cA_4_ degradation, as viral proteins are often multifunctional, a property that is favored when the alternate activities are complementary^51^ or potentially expand the range of action of the protein to different scenarios (e.g. presence/absence of CRISPR-Cas targeting, Fig. 5).

**Figure 5.**
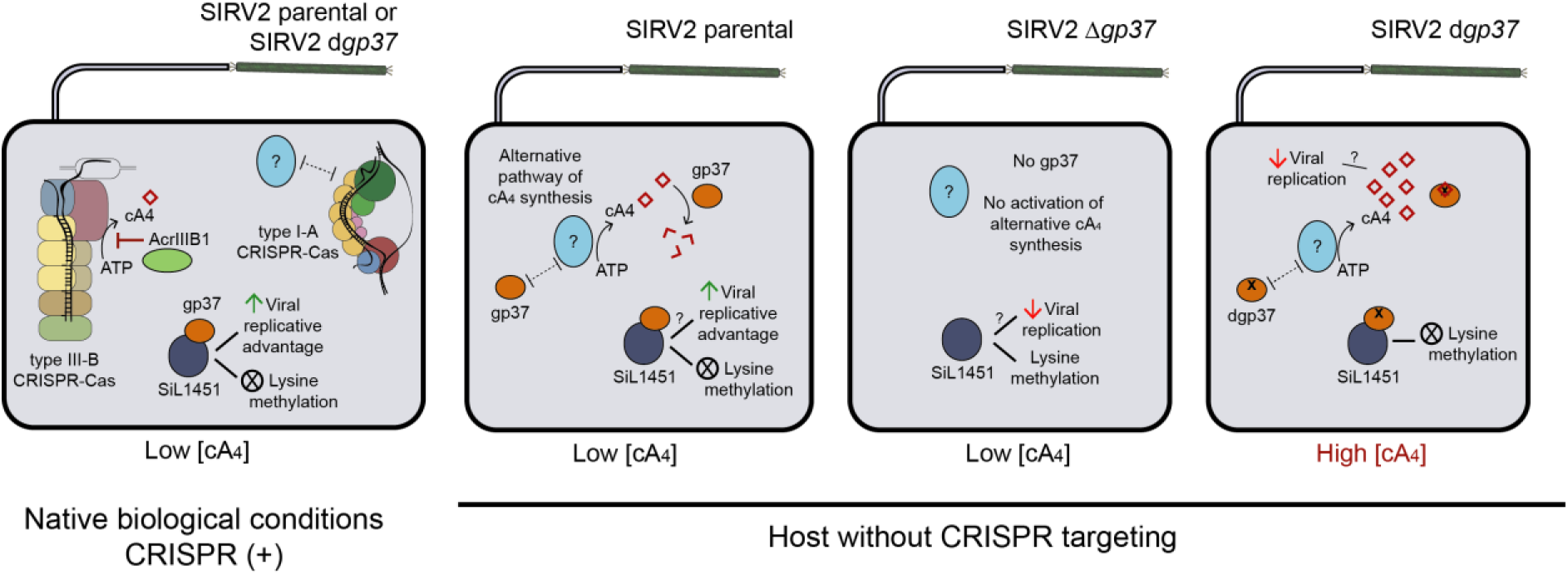
Model of the effect of multifunctional gp37 during SIRV2 infection under diverse scenarios. *First panel*: During SIRV2-infection of the native host *S. islandicus* LAL14/1 intracellular cA4 levels are kept low through the inhibition by viral AcrIIIB1 of the type III-B CRISPR-Cas and by the effect of the activation of type I-A CRISPR-Cas over alternative sources of cA4 production. Thus, the RING nuclease activity of gp37 is dispensable in this scenario. Interaction between gp37 and lysine methyltransferase SiL_1451 results in the inhibition of the latter, abrogation of cellular protein lysine methylation and a positive effect on viral replication through a yet unknown mechanism. Second to fourth panels correspond to infection with the indicated SIRV2 strains of a host without CRISPR-Cas mediated targeting of the virus. *Second panel*: Infection by parental SIRV2. Lack of I-A CRISPR-Cas targeting and presence of gp37 activates an alternate cA4 synthesis pathway unrelated to type III-B CRISPR-Cas. This cA4 is rapidly degraded by the RING nuclease activity of gp37, keeping cA4 intracellular levels low. Similar to the first panel, inhibition of SiL_1451 by gp37 is beneficial for viral replication. *Third panel*: Infection by SIRV2 Δgp37 mutant lacking gp37. Absence of gp37 fails to trigger the alternate cA4 synthesis pathway, thus intracellular concentration of this nucleotide is low and RING nuclease activity is not necessary. Nevertheless, active SiL_1451 has a mild negative impact on SIRV2 replication through an unidentified mechanism. *Fourth panel*: Infection by SIRV2 d gp37 encoding a gp37 with abrogated RING nuclease activity. Presence of gp37 activates alternate cA4 production, which negatively affects viral replication, overcoming the positive impact of the gp37-mediated inhibition of SiL_1451. Dotted lines represent a yet unidentified direct or indirect interaction between two protein complexes. Interrogation signs indicate unknown mechanism of action.

Equally elusive is the role of the lysine methyltransferase SiL_1451, mediating ample protein methylation in crenarchaea whose functional significance has not been elucidated^38^. Our work establishes a link between SiL_1451 and cellular defense and/or stress response. Interestingly, a knockout mutant of the lysine methyltransferase shows differential expression of genes involved in metabolism and genes without functional annotation, plus several genes located in the variable region of the *S. islandicus* strain probably corresponding to integrated viruses^38^ (Fig. S9), supporting a role of protein methylation in stress response. Additionally, differential methylation in response to heat-shock of Cren7, a SiL_1451 substrate, affects the DNA-binding ability of Cren7 potentially impacting chromosomal regulation, suggesting a role of SiL_1451 in mediating epigenetic regulation in response to environmental stress^52^. The mechanism by which inhibition of SiL_1451 methyltransferase activity aids SIRV2 replication remains to be elucidated. The replicative advantage conferred by gp37 is an example about how mild effects become relevant at the population level and can mediate the selective sweep of a strain in the population.

The use of heterologous systems have allowed the discovery and characterization of multiple novel defense systems in prokaryotes^1–4,21–24^ and constitute an essential tool when a cultivable isolate of the system under study is not available or it lacks tools for its genetic manipulation. However, heterologous systems may introduce artifacts that lead to erroneous conclusions, among other reasons due to changes in expression level, location or timing with respect to the native system. Therefore, it is of uttermost importance to characterize novel proteins under their native biological conditions whenever possible.

## Acknowledgements

We are truly grateful to Jinzhong Lin for providing purified *S. islandicus* Cmr-α effector complex and SS1-46 target RNA. This work was supported by the Danish Council for Independent Research/Natural Sciences [DFF-0135-00402] and Novo Nordisk Foundation/Hallas Møller Ascending Investigator Grant [NNF17OC0031154] to X.P.

## Author contributions

L.M-A. designed and performed experiments, oversaw the work, analyzed the data and wrote the manuscript. D.S. generated viral strains and performed one-step growth curves. A.S-D. purified proteins, carried out cA4-cleavage assays and contributed to the competition assays. Y.B-C. generated the virus mutant SIRV2 *P_ara_-gp29*. X.P. conceived the project, oversaw the work and edited the manuscript. All authors contributed to data analysis.

## FIGURES

**Figure S1.**
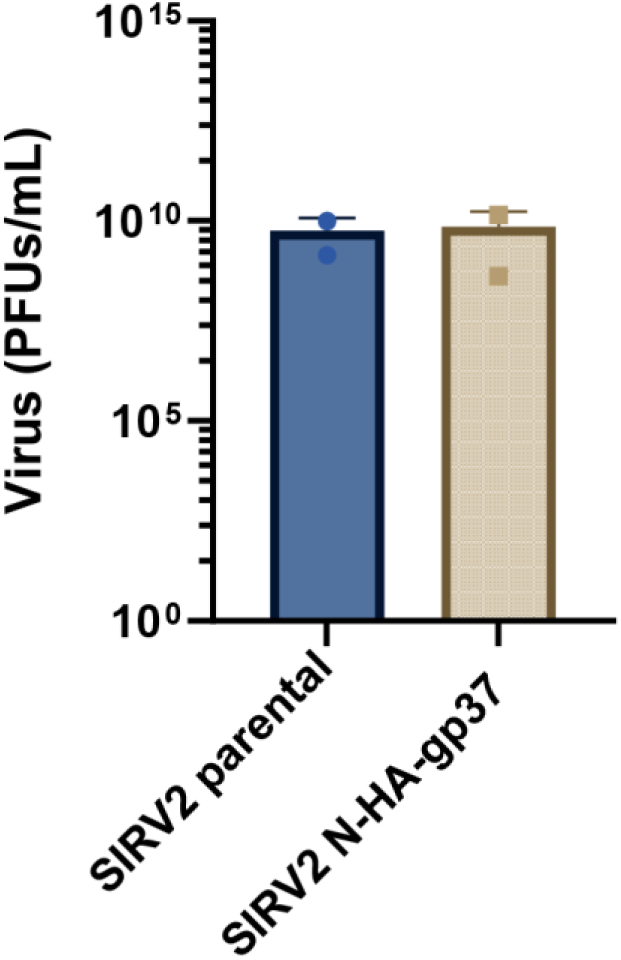
Virus mutant SIRV2 N-HA-gp37 with an N-terminally hemagglutinin tagged gp37 is similarly infective as the parental virus. CRISPR (-) cultures were infected with the corresponding virus at an MOI of 0.01 and virus in the supernatant was determined after 36 h.

**Figure S2.**
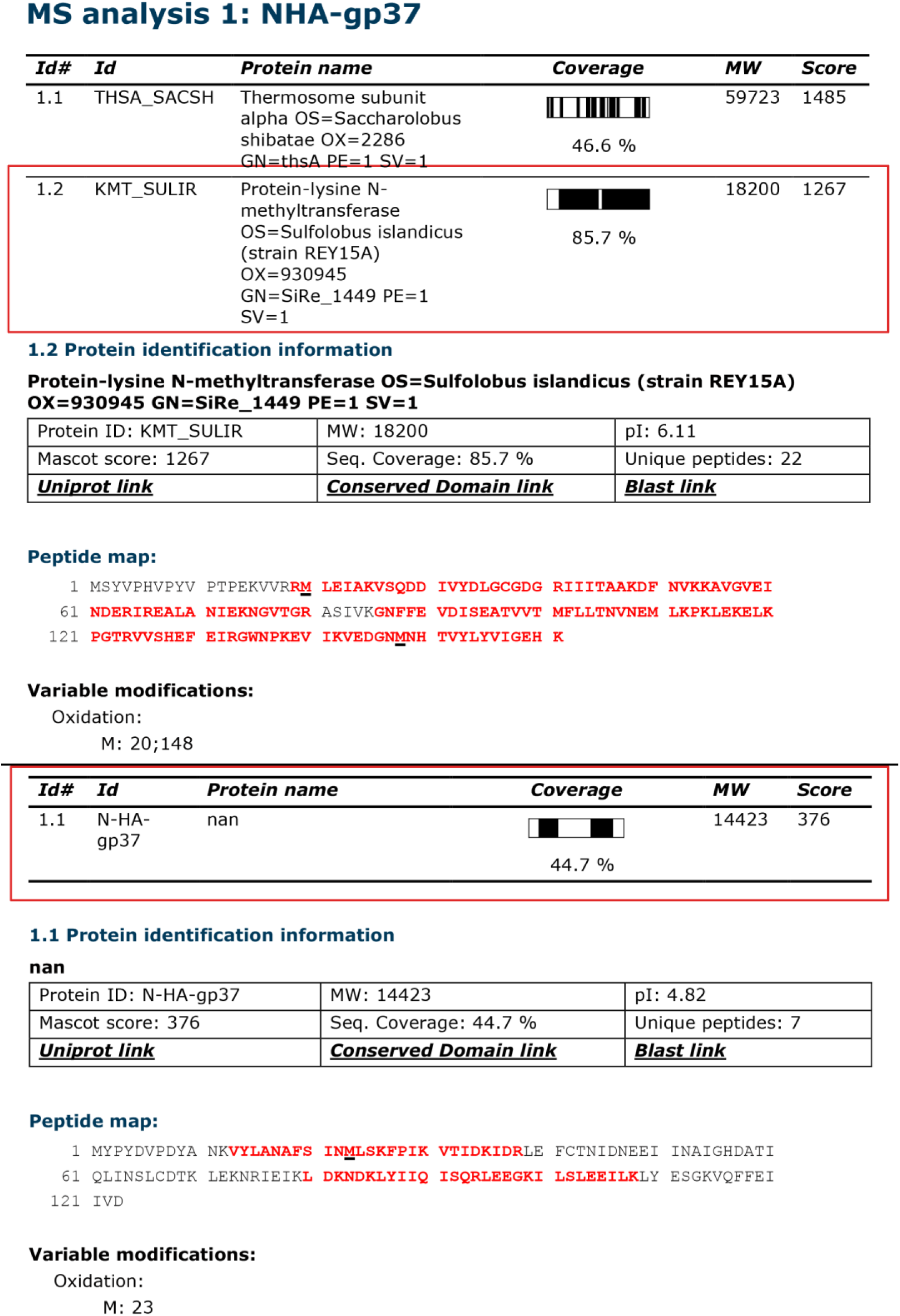
Nanoscale liquid chromatography coupled to mass spectrometry results of protein eluate in Fig. 3A. The analysis identifies the 18 kDa band as the lysine methyltransferase SiRe_1449 corresponding to SiL_1451 in *S. islandicus* LAL14/1 and the 14.4 kDa band as SIRV2 gp37.

**Figure S3.**
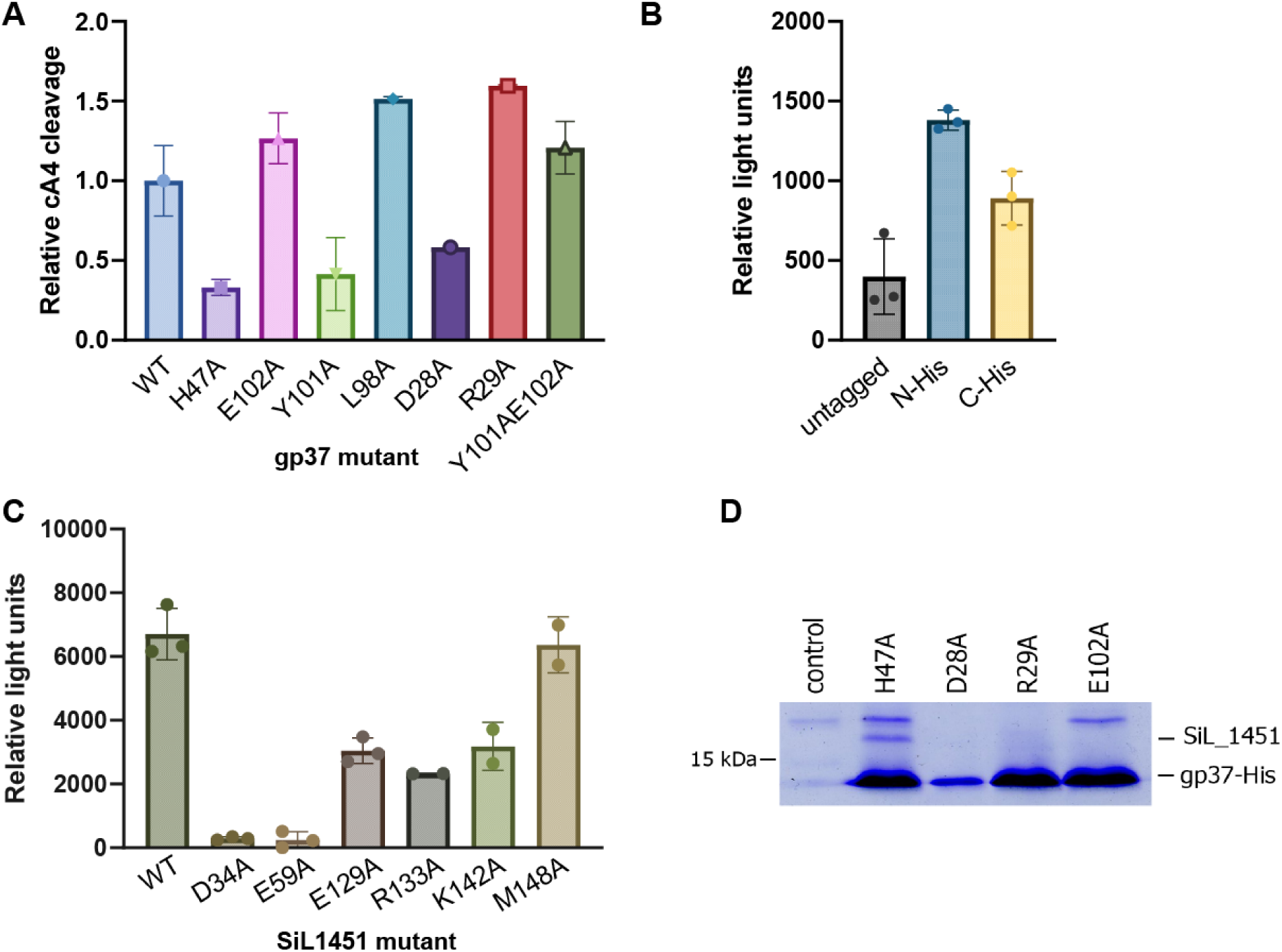
Catalytic activity of gp37 and SiL_1451 alanine mutants of putative gp37-SiL_1451 interaction residues. **A**) cA_4_ cleavage activity of gp37 mutants relative to the wild-type (WT) protein. Mutant H47A is a catalytically defective RING nuclease, as previously described by Athukoralage et al. (2019). Mutants Y101A and D28A show abrogated cA_4_ cleavage capacity and were not used in downstream assays. **B)** Effect of His-tag placement on the methyltransferase activity of SiL_1451. Methyltransferase activity was measured with the MTase-Glo™ assay using Cren7 as the substrate, as described in methods. The untagged SiL_1451 protein was obtained from enterokinase treatment of the N-terminally His-tagged recombinant protein. The N-terminally tagged SiL_1451 (N-His) was used for mutant construction as it displays the highest methyltransferase activity. **C)** Methyltransferase activity of SiL_1451 mutants. Mutants D28A and E59A are unable to bind the cofactor SAM as reported previously (Chu et al. 2010) and thus display no methyltransferase activity. All mutants tested, with the exception of M148A, have reduced methyltransferase activity. The data correspond to the average of two or three biological replicates. Bars correspond to the standard deviation. **D)** Replicate of the C-terminally his-tagged gp37 pull-down experiment of Fig. 3F.

**Figure S4.**
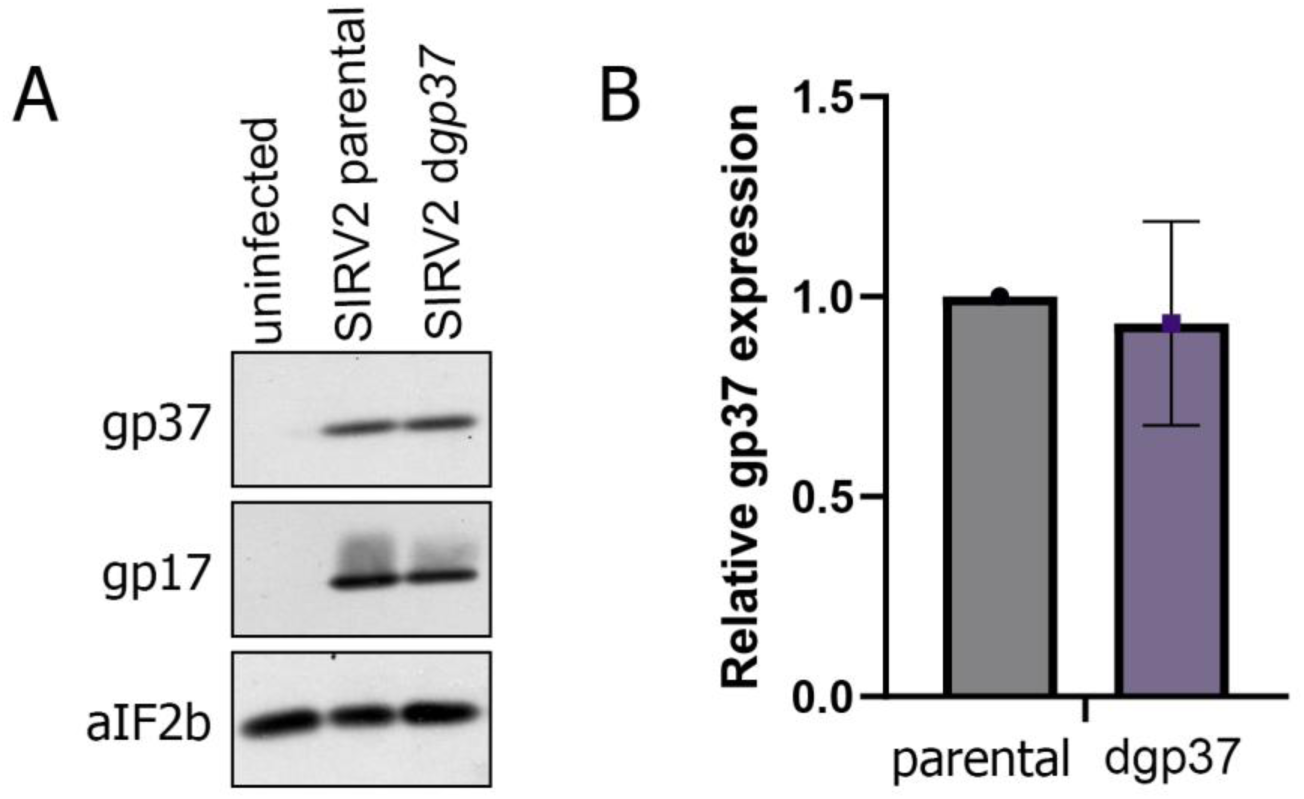
Expression level of gp37 in the virus mutant SIRV2 dgp37 is comparable to the expression in the parental strain. CRISPR (-) cultures were infected with an MOI of 3 with the parental strain SIRV2 or mutant SIRV2 d gp37 encoding a catalytically dead gp37 (H47A mutant). Samples were collected at 3 hpi and analyzed by western blot. Data was normalized using aIF2β as loading control and dgp37 protein level is reported relative to wild-type gp37 expression. **A)** Representative western blot of infected cellular lysates (*n*=3). **B)** Relative average d gp37 expression with s.d. from three biological replicates.

**Figure S5.**
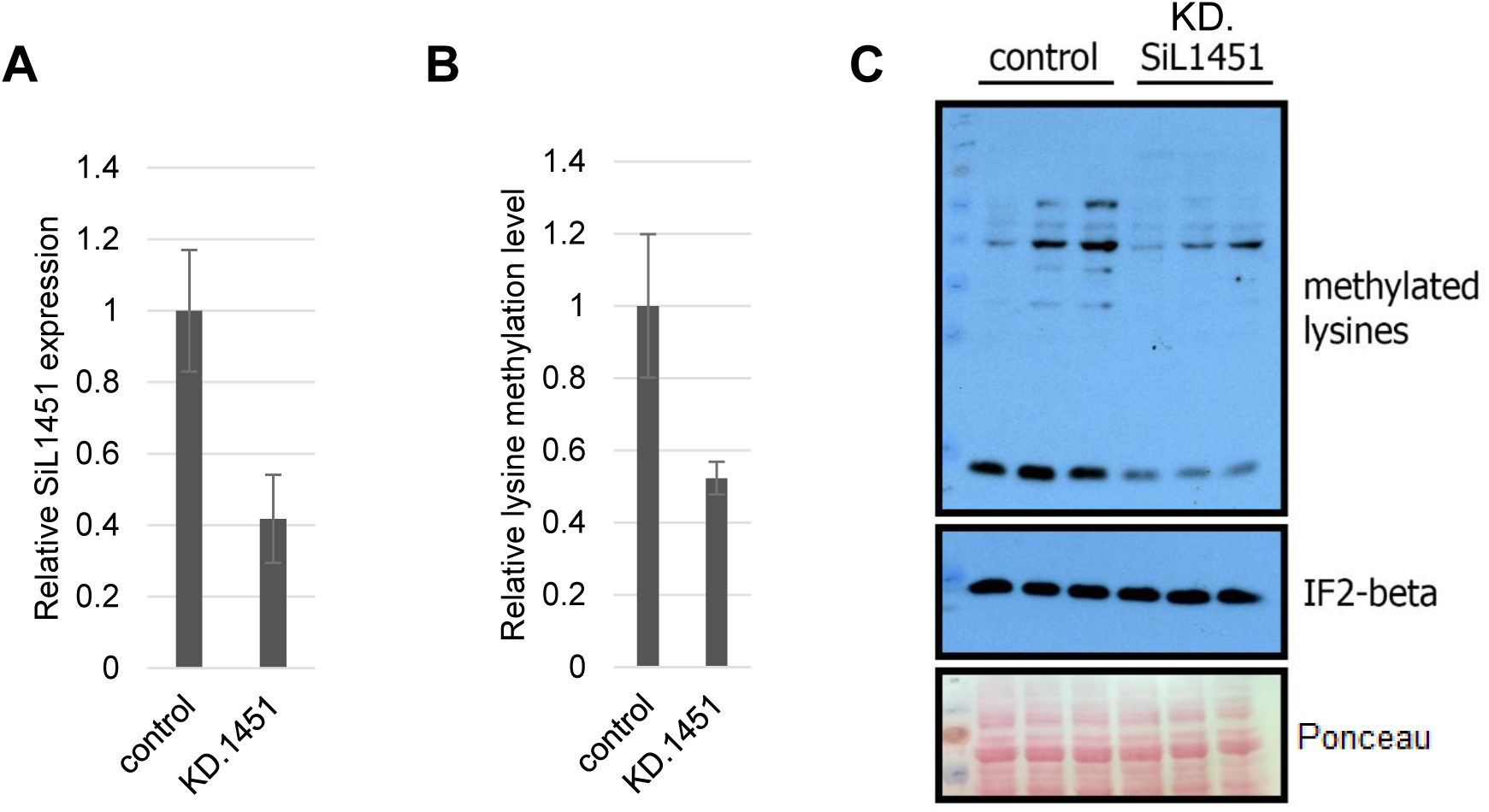
CRISPR-based silencing of SiL_1451 expression. **A**) Transcription level of *siL_1451* determined by RT-qPCR in samples of control culture (harboring an empty plasmid) and SiL_1451 knock-down strain (KD.1451) at OD600 = 0.2. Data represent the average and s.d. *siL_1451* mRNA of three biological replicates. **B)** Relative lysine methylation level of cultures in panel A as determined by western blot. Data correspond to average lysine methylation and s.d. of three biological replicates. **C)** Representative western blot of cellular lysates in panel B. IF2β and Ponceau-stained total protein were used as loading control.

**Figure S6.**
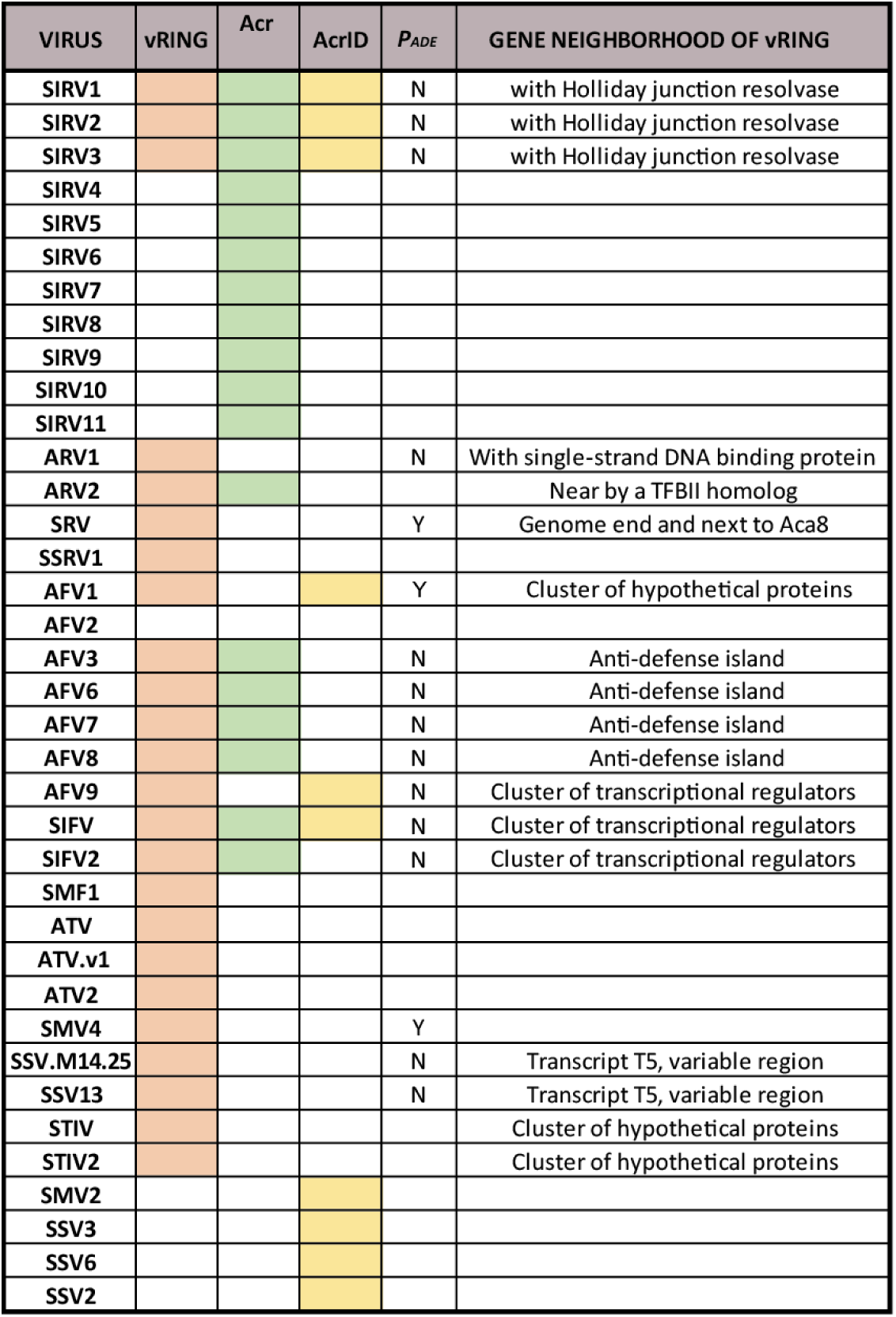
Genome neighborhood analysis of vRING nuclease homologs in archaeal viruses. Viruses with homologs of vRING nuclease proteins, AcrIIIB-1 or AcrID were retrieved and the location of vRING nuclease homologs was analyzed. vRING homologs under the control of the anti-defense regulatory sequence (*P_ADE_*) are indicated, while homologs in the vicinity of an Acr or Aca protein and in a cluster of small hypothetical genes coded in the same strand were annotated as encoded in an anti-defense island. Transcript T5 refers to one of the transcriptional units of fuselloviruses that typically encodes variable genes (Aulitto et al. 2023).

**Figure S7.**
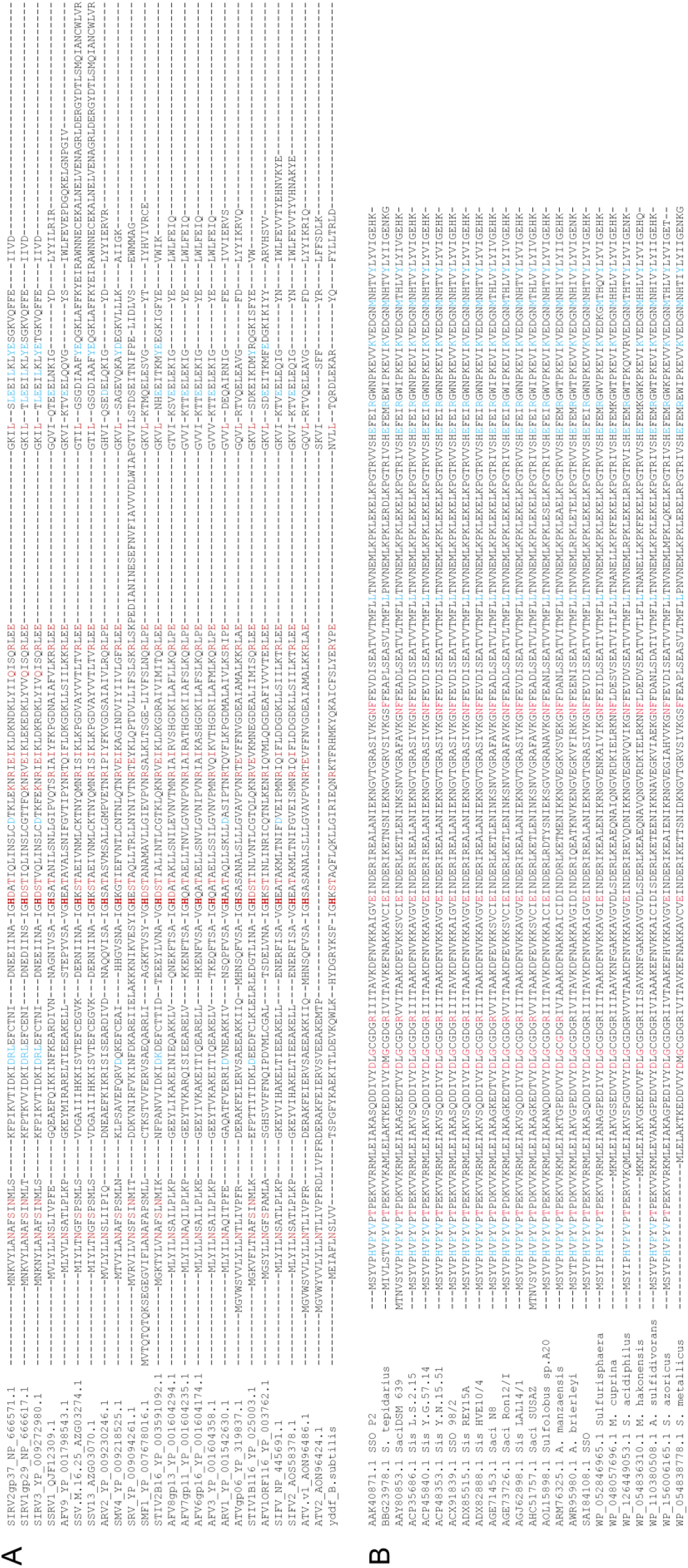
Conservation of crenarchaeal vRING and lysine methyltransferase homologs. **A**) Alignment of vRING nuclease proteins in crenarchaeal viruses. **B)** Alignment of lysine methyltransferase homologs in the Sulfolobales. Aminoacids marked in red are important for cA_4_ cleavage (A) or SAM binding (B) (Athukoralage et al. 2019, Chu et al. 2016). Residues predicted to be involved in the gp37-SiL_1451 interaction are marked in blue.

**Figure S8.**
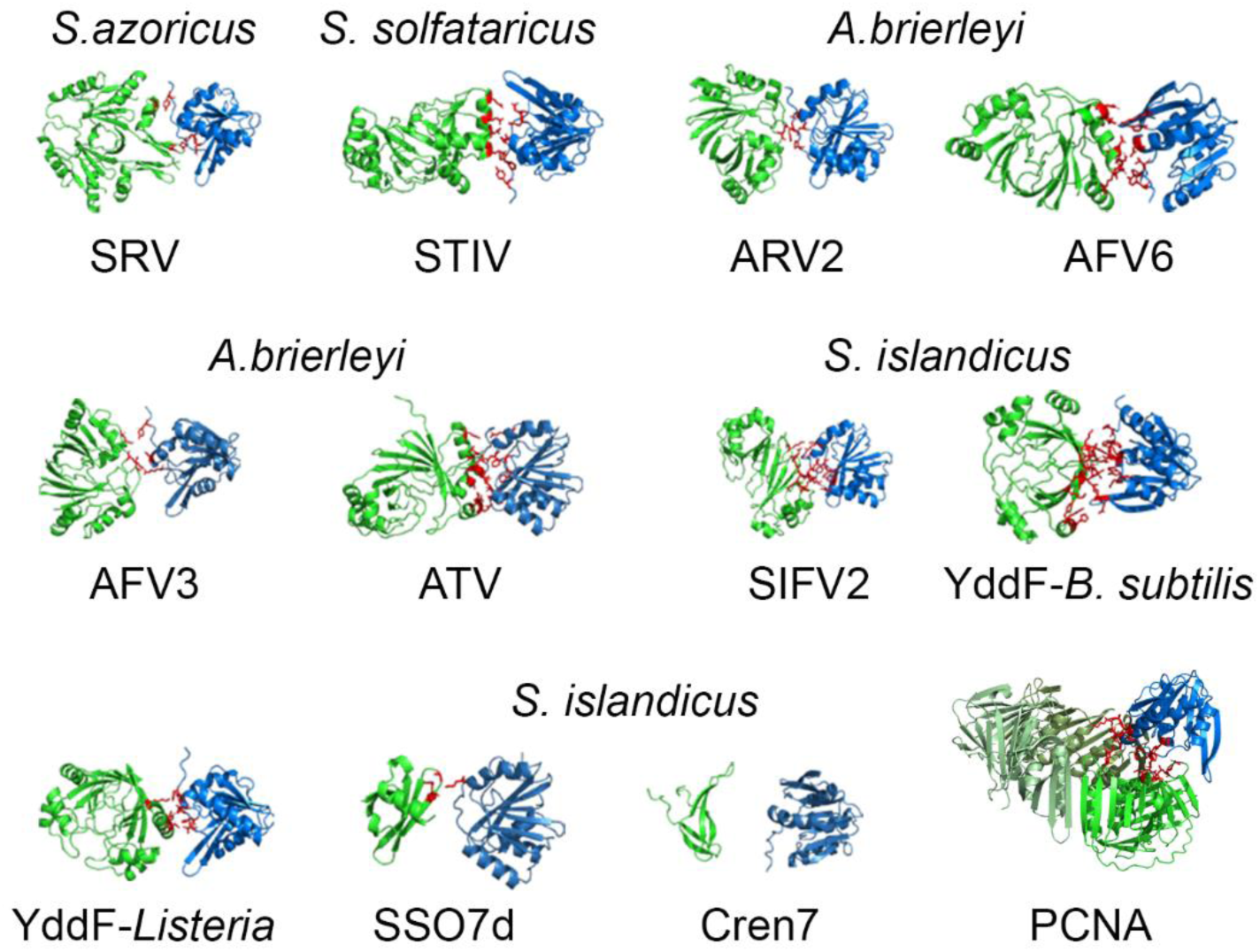
AlphaFold2 models of vRING nuclease proteins and their cognate lysine methyltransferase. The vRING homolog is colored in green and the lysine methyltransferase is indicated in blue. Putative interacting residues are colored in red.

**Figure S9.**
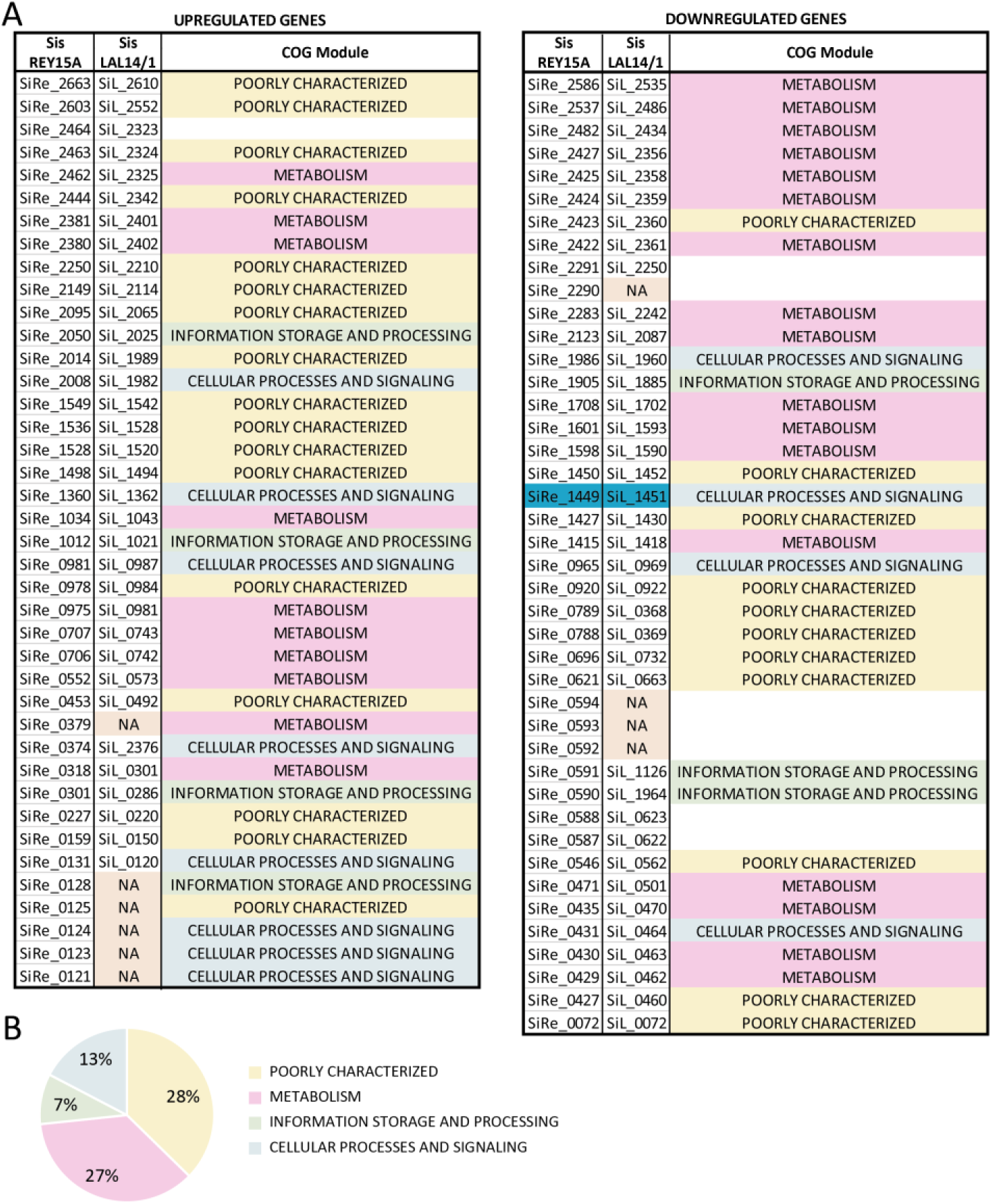
Differentially regulated genes in a lysine methyltransferase knock-out strain. **A**) List of differentially regulated genes in the strain *S. islandicus* REY15A Δ*sil_1451*, their corresponding homolog in *S. islandicus* LAL14/1 and their COG Module annotation. Data adapted from Chu et al. 2016. **B)** Pie chart of the COG Module functional annotation of genes in panel A.

